# MT-125 Inhibits Non-Muscle Myosin IIA and IIB, Synergizes with Oncogenic Kinase Inhibitors, and Prolongs Survival in Glioblastoma

**DOI:** 10.1101/2024.04.27.591399

**Authors:** Rajappa Kenchappa, Laszlo Radnai, Erica J. Young, Natanael Zarco, Li Lin, Athanassios Dovas, Christian T. Meyer, Ashley Haddock, Alice Hall, Peter Canoll, Michael D. Cameron, Naveen KH Nagaiah, Gavin Rumbaugh, Patrick R. Griffin, Theodore M. Kamenecka, Courtney A. Miller, Steven S. Rosenfeld

## Abstract

We have identified a NMIIA and IIB-specific small molecule inhibitor, MT-125, and have studied its effects in GBM. MT-125 has high brain penetrance and retention and an excellent safety profile; blocks GBM invasion and cytokinesis, consistent with the known roles of NMII; and prolongs survival as a single agent in murine GBM models. MT-125 increases signaling along both the PDGFR- and MAPK-driven pathways through a mechanism that involves the upregulation of reactive oxygen species, and it synergizes with FDA-approved PDGFR and mTOR inhibitors *in vitro*. Combining MT-125 with sunitinib, a PDGFR inhibitor, or paxalisib, a combined PI3 Kinase/mTOR inhibitor significantly improves survival in orthotopic GBM models over either drug alone, and in the case of sunitinib, markedly prolongs survival in ∼40% of mice. Our results provide a powerful rationale for developing NMII targeting strategies to treat cancer and demonstrate that MT-125 has strong clinical potential for the treatment of GBM.

**Highlights:**

- MT-125 is a highly specific small molecule inhibitor of non-muscle myosin IIA and IIB, is well-tolerated, and achieves therapeutic concentrations in the brain with systemic dosing.
- Treating preclinical models of glioblastoma with MT-125 produces durable improvements in survival.
- MT-125 stimulates PDGFR- and MAPK-driven signaling in glioblastoma and increases dependency on these pathways.
- Combining MT-125 with an FDA-approved PDGFR inhibitor in a mouse GBM model synergizes to improve median survival over either drug alone, and produces tumor free, prolonged survival in over 40% of mice.

## INTRODUCTION

The myosins are a superfamily of molecular motors (Odronitz and Kollmar, 2007) and are widely expressed. Among these are members of the myosin II class, which includes skeletal, smooth, and cardiac myosin II along with three non-muscle myosin II (NMII) paralogs, referred to as NMIIA, IIB, and IIC. NMII plays multiple roles in cell physiology, including motility, cytokinesis, mitochondrial fission, and signaling (Straight, *et al.,* 2003; Garrido-Casado, *et al*., 2021; Quintanilla, Hammer, and Beach, 2023; Najafabadi, *et al.,* 2022; Stricker, *et al*., 2013, Kim *et al.,* 2012; Zheng et al., 2021). However, these NMII-driven functions are also important in supporting pathological states. An example of this can be seen in glioblastoma (GBM), the most common and malignant of primary brain tumors (Wen, *et al.,* 2020). Aberrant signaling through activating mutations/amplifications of oncogenic kinases are frequently found in GBM (Szerlip *et al*. 2012). Nevertheless, clinical trials of oncogenic kinase inhibitors have been disappointing (Bai *et al.,* 2011; Mellinghoff, *et al.,* 2012). This likely reflects a redundancy in signaling pathways, allowing cells to circumvent blockade of one pathway by activating another. NMII paralogs are downstream of many of these oncogenic kinases at points where signaling pathways converge. This implies that direct inhibition of NMII could be effective at blocking tumor physiology, even with multiple activated upstream oncogenic effectors. In support of this, we found that direct inhibition of NMII with blebbistatin, a small molecule inhibitor of both muscle and non-muscle myosin II, could block glioma dispersion, even when both PDGFR and EGFR-driven pathways were simultaneously activated (Ivkovic *et al*., 2012). More recently, we also showed that co-deleting both NMIIA and IIB prevents tumorigenesis in genetically engineered mouse models (GEMMs) of GBM (Picariello *et al*., 2019). This study also found that deleting just NMIIA activates SRC, ERK1/2, YAP, and NFkB (Picariello *et al*., 2019), consistent with earlier reports that NMIIA could function as a tumor suppressor (Coaxum et al., 2017; Conti et al., 2015; Schramek et al., 2014). Taken together, these findings suggest that while a drug that inhibits both NMIIA and IIB might increase oncogenic signaling, through its inhibition of NMIIA, such a drug would nevertheless prevent this upregulated signaling from translating into a more aggressive tumor, since inhibiting both NMIIA and IIB together should block kinase driven proliferation and invasion. Furthermore, increasing oncogenic signaling might induce *oncogene addiction* (Weinstein & Joe, 2008) to these pathways, leading to enhanced tumor kill by combining an NMIIA/IIB inhibitor with oncogenic kinase inhibitors.

These findings argue that a dual inhibitor selective for NMIIA and NMIIB might be highly active against GBM and provide a new avenue for therapeutic applications. However, capitalizing on this information might be limited by the possibility that an NMII inhibitor could be toxic, due to the wide distribution of NMII paralogs in normal tissues (Sellers and Heissler, 2019). In this study, we describe the development and excellent safety profile of MT-125, a dual NMIIA/IIB small molecule inhibitor, and examine its mechanism of action and efficacy in murine models of GBM.

## RESULTS

### MT-125 is a small molecule inhibitor that is based on the structure of blebbistatin and is highly selective for NMIIA and IIB

We evaluated a library of derivatives based on the pan-myosin II small molecule inhibitor blebbistatin, and identified MT-125 for its nearly equal potency against NMIIA (K_i,NMIIA_ = 2.7 ± 0.2 µM) and NMIIB (EC_50_ = 1.7 ± 0.1 μM) (**Fig. 1A-B**). MT-125 is not cytotoxic at any dose using a COS7 cell-based assay (**Fig. 1B***, red*). We designed MT-125 to provide ∼20-30-fold selectivity for NMIIA and IIB over cardiac myosin II (K_I,CMII_ = 50 ± 10 µM, **Fig. 1A**). Based on our prior work, we expected this extremely low CMII potency would translate into excellent *in vivo* tolerability. We have reported MT-134, a skeletal myosin II-specific derivative of blebbistatin with low CMII inhibition (K_I,CMII_ = 83 µM) is well-tolerated *in vivo* up to the highest dose tested (10 mg/kg; Radnai *et al*, 2021).

**Figure 1:**
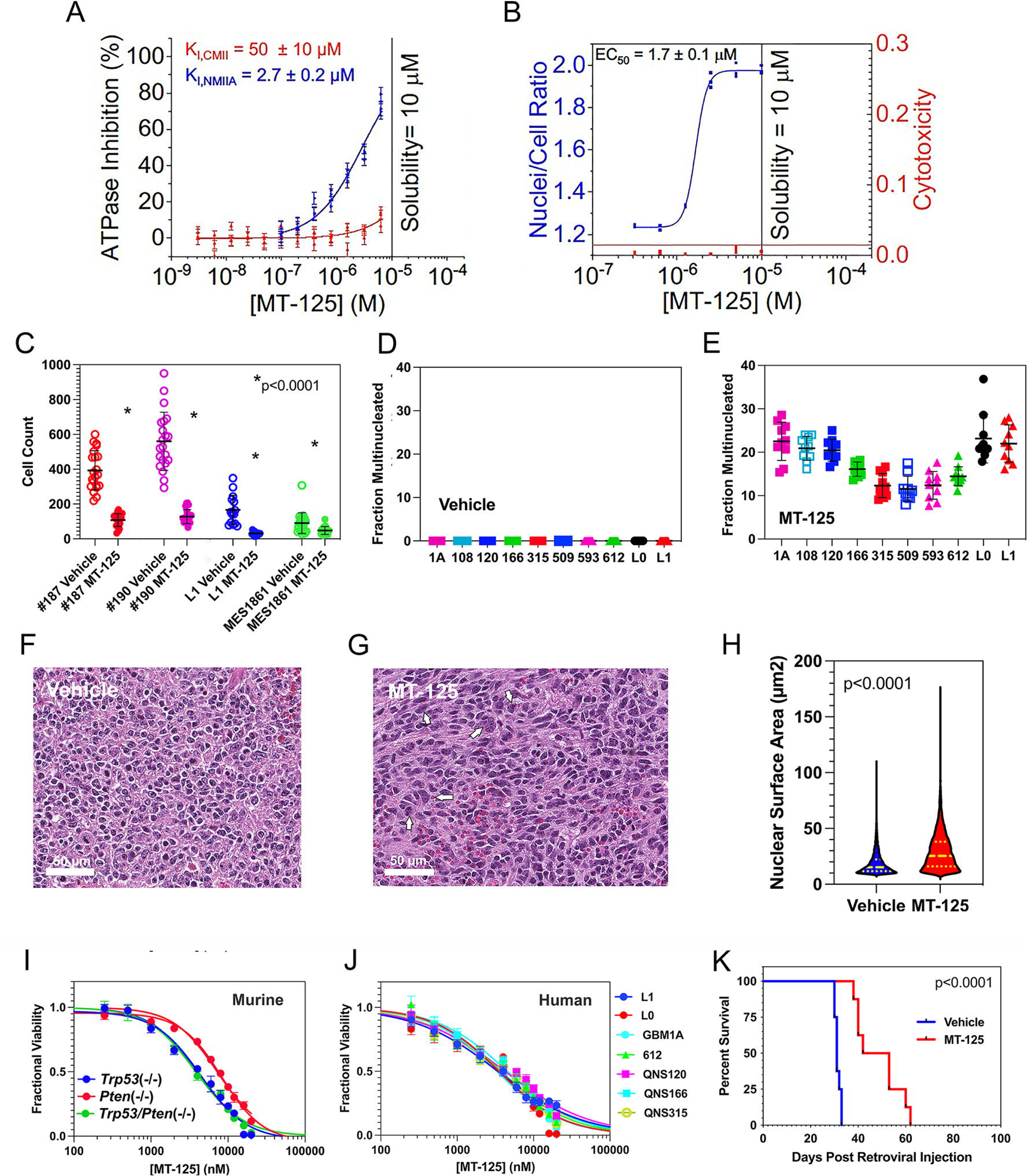
MT-125 phenocopies the effects of NMII deletion and is therapeutic in a murine model of GBM. **(A)** The potencies of MT-125 for CMII and NMIIA were determined in an NADH-linked ATPase assay (PMID: 31475972). **(B)** A COS7-cell based cytokinesis assay was used to assess potency for NMIIB (PMID: 33204739; PMID: 15774463). **(C).** Invasion of three human (187, 190, L1) and one murine GBM line (MES1861) through 3 µm Transwell membranes in the presence of vehicle (*open*) or 5 µM MT-125 (*closed*). **(D** & **E).** Ten human GBM lines were treated with vehicle (**B**) or MT-125 (**C**) for 72 hours, and the fraction of multinucleated cells were measured in 8 replicates. (**F-H**). *Trp53*(−/−) GEMMs were treated with vehicle (**F**) or MT-125 daily (**G**) for 14 days. MT-125 treatment induces multinucleation (**G**, *arrows*) and increases mean nuclear cross-sectional area by 53% (**H***, p<0.0001*). (**I,J**). Dose responses for three murine (**I**) and seven human (**J**) GBM lines to MT-125. Values of EC_50_ and Hill coefficient are summarized in **Table S2**). (**K**). Kaplan Meier survival curves for *Trp53*(−/−) GEMMs treated with vehicle (DMSO, *blue*) or MT-125 (*red*).

### Pharmacokinetic, metabolism, and safety evaluation of MT-125

We performed absorption, distribution, metabolism, and excretion (ADME) profiling to determine the drug-like properties of MT-125. *In vitro* absorption studies of MT-125 at a drug concentration of 10 μM demonstrated little effect on passive permeability. Inhibition of BSEP, MATE1 and MATE-2 transporters occurred in a range of 56-86%, with minimal activity in the Caco-2 A-B and B-A permeability assays. In intrinsic clearance assays using liver microsomes from human, dog, rat, and mouse, the half-life of MT-125 ranged from 5.8 to 40.2 minutes (human < mouse < dog < rat). In intrinsic clearance assays using cryopreserved hepatocytes, the half-life ranged from 48.1 to 113.6 minutes (mouse < rat < human < dog). These data suggested a relatively short half-life for MT-125 *in vivo*.

To identify enzymes responsible for MT-125 metabolism, cytochrome P450 (CYP) phenotyping and induction was performed. In CYP intrinsic clearance assays, the half-life for most P450s was greater than 60 minutes, except for CYP1A2 (39.8 mins) and CYP2C8 (32.6 mins), suggesting these two P450s metabolize MT-125. MT-125 did not demonstrate time-dependent inhibition at a concentration of 10 μM and was below the 37% inhibition displayed by the reference compounds for all CYP450s tested. MT-125 *in vitro* induces of CYP1A2 and CYP2B6 at dose of 10 μM and higher, but not at 1 μM, and induces CYP3A4 at 1 μM in 2 of 3 donors, and in all donors at 10 μM, suggesting a potential for drug-drug interactions that would need to be considered. However, this risk is minimal at lower doses. We found that MT-125 is not a P-gp substrate, suggesting it should have good brain retention. hERG-CHO patch clamp and an ion channel activity panel were assessed as a measure of cardiac safety *in vitro*. An IC_50_ for MT-125 could not be calculated for Nav1.5, Kv4.3/KChiP2, hERG, KCNQ1/mink, and Kir2.1, as inhibition greater than 25% was not observed. These results in combination with the lack of CMII inhibition indicate that MT-125 should not impact cardiac function.

We also screened 78 G-protein coupled receptors (GPCRs), enzymes, ion channels, transporters, and nuclear receptors to look for off-target activity. For 61 of these, IC/EC_50_’s could not be calculated due to very low activity. For the 17 other targets the IC/EC_50_’s were in the range of 1-5 μM. MT-125 has an IC_50_ for hERG of 5.4 μM, with a maximum response of 65% of the control. However, in the hERG-CHO electrophysiology assay, MT-125 has an estimated IC_50_ greater than 10 μM. Cav1.2 was consistent between the two profling screens with an estimated IC_50_ greater than 10 μM. The remaining 15 targets were GPCRs involved in cAMP signaling and MT-125 displays both agonist and antagonist activity. Taken together, MT-125 displays an acceptable selectivity profile for drug development advancement.

*In vivo* pharmacokinetic profiling of MT-125 was performed with several routes of systemic administration to establish the optimal route of administration, the degree of brain penetrance and retention, the general pharmacokinetic properties, and initial information on tolerability. At 2 mg/kg (IV), MT-125 had a maximal concentration (C_max_) of 24.5 ± 3.2 µM, AUC_last_ of 12.2 ± 2.0 µM.hr, and T_1/2_ of 2.3 hr in plasma and was well-tolerated with no adverse clinical signs. Oral dosing at 5 times the IV dose (10 mg/kg, PO) was also well-tolerated, but demonstrated fairly low oral bioavailability (F% = 12.0), with a C_max_ of 18.8 ± 1.5 µM, AUC_last_ of 7.3 ± 2.2 µM.hr, and T_1/2_ of 2.0 hours. Due to solubility limitations, higher doses resulted in a suspension, which showed poorer absorption. Therefore, PO dosing was not pursued. Subcutaneous (SC) dosing, a route of administration that would be preferred for this patient population over IV, at 5 and 10 mg/kg was also well-tolerated and showed excellent systemic exposure (**Table S1**). Importantly, very high brain levels were achieved rapidly after administration. For instance, at 5 mg/kg (SC), maximal concentrations of 9.1 ± 1.0 µM (C_max_) were achieved in the brain within 10 minutes of injection (T_max_) and were double that present in plasma at the same time point (B:P = 2.0). Further, MT-125 remained at measureable levels in the brain well after injection, resulting in brain AUC_last_ of 5.4 ± 0.8 µM.hr and a T_1/2_ of 10.5 hours (**Table S1**). The brain to plasma ratio (B:P) at 5 mg/kg was 2.4.

Given that several of the *in vivo* efficacy experiments in this study employed intraperitoneal (IP) administration, we also compared SC and IP dosing at 10 mg/kg to determine relative differences in the plasma and brain drug levels via the different routes of administration within 30 minutes of injection. Markedly lower plasma concentrations of MT-125 were observed 30 mins after IP and SC administration (IP = 0.7 ± 0.3 µM, SC = 2.5 ± 0.2 µM). However, because only a single time point (30 minutes) was assessed after compound administration, it is worth noting these are not the C_max_ values, and the kinetics are likely different between the two routes of administration. As with SC administration, excellent brain penetration was observed with IP administration. Thirty minutes after administering 10 mg/kg IP, the brain concentration was greater than that found in plasma (brain = 1.0 ± 0.3 µM, with a brain to plasma ratio of 1.6).

### MT-125 demonstrates low toxicity vivo and has a very favorable therapeutic index

Two weeks of daily MT-125 adminstration (10 mg/kg, SC) was well-tolerated in mice, with no change in body weight (Day x Treatment: F_(13, 91)_ = 1.5, *p* > 0.05; **Fig. S1A**). Moreover, there was no change in clinical chemistry or hematology values after 14 days of MT-125 administration, compared to vehicle (For all measures *p* > 0.05, **Fig. S1B**). To determine the maximum tolerated dose (MTD) for a single administration of MT-125, rats were given one injection of MT-125 at 60, 70 or 90 mg/kg (SC). Blood samples were collected one hour later for toxicokinetic analysis and twenty-four hours later for hematology and clinical chemistry. There were no observable adverse clinical effects or changes in body weight at any dose tested (Day x Treatment: F_(2, 6)_ = 1.1, *p* > 0.05; **Fig. S2A**). Plasma concentrations one hour post-dose indicated that MT-125 was absorbed (60 mg/kg: 4.8 ± 0.6 µM, 70 mg/kg: 3.6 ± 0.4 µM, 90 mg/kg: 4.4 ± 0.3 µM). However, the similarity of plasma concentrations across the dose levels suggests that 60 mg/kg may contain the maximum amount of compound that can be absorbed at a single SC injection site. All clinical chemistry and hematology values were within the normal range 24 hours after dosing (**Fig. S2B**).

We next injected rats with vehicle, 20 mg/kg, or 40 mg/kg (SC) MT-125 daily for 7 days. Vehicle and MT-125 at 20 mg/kg produced no drug-related adverse health effects or effects on normal age-related body weight gain (during dosing: Veh vs 20 mg/kg, *p* > 0.05; during the 7-day recovery period: Veh vs 20 mg/kg, *p* > 0.05; **Fig. S3A**). Repeated injection of 40 mg/kg in males led to slight, temporary changes in breathing, as well as drooling that developed 2 to 4 hours post-dose and resolved within 30 minutes to one hour. Male rats treated with 40 mg/kg MT-125 did not gain or lose weight during the dosing period. However, they did gain weight during the recovery period (40 mg/kg, males: Day 1 vs Day 8, *p* > 0.05; during dosing: Veh vs 40 mg/kg, *p* < 0.0001; during the 7-day recovery period: Veh vs 40 mg/kg, *p* > 0.05; **Fig. S3A**). Females exhibited no adverse health effects or changes in body weight (Day 1 vs Day 8: *p* > 0.05, **Fig. S3A**). All clinical chemistry and hematology values in males and females in the vehicle control and at both dose levels were within normal ranges at the end of 7 days of treatment and following a 7-day recovery period (**Fig. S3B**). Toxicokinetics were evaluated across a 6-point timecourse (30 minutes to 24 hours post-dose) on Days 1 and 7, plus a pre-dose collection on Day 7. The expected dose-dependent increases in plasma concentrations were observed, with no sex differences. At the 40 mg/kg dose level, AUC_last_ for males was 20.4 ± 2.1 µM.hr and females was 17.0 ± 1.7 µM.hr (p > 0.05). A slight accumulation was seen over the 7 days of administration, as plasma levels prior to the Day 7 dose administration were 0.43 µM ± 0.05 µM (**Fig. S3C**). Based on these results the STD 10 (10% mortality over the study duration), which is used to benchmark toxicology limits for oncology therapeutics in rodents, is >40 mg/kg.

### MT-125 phenocopies the effects of NMII deletion on GBM invasion and proliferation, and prolongs survival in a murine GBM model

Deletion of NMIIA and NMIIB impairs proliferation, and targeting NMII with blebbistatin blocks invasion (Beadle et al., 2008). To confirm that MT-125 works similarly, we measured *in vitro* invasion across 3 µm Transwell pores after 8 hours of exposure to vehicle (DMSO) or 5 µM MT-125 for one murine (MES1861) and three human (187, 190, L1) GBM cell lines (**Fig. 1C**). In each case, MT-125 significantly impairs Transwell invasion (*p<0.0001*). NMII is required for cytokinesis, and its inhibition leads to multinucleation and aneuploidy (Newell-Litwa, *et al.,* 2015; Najafabadi, *et al.,* 2022). We found that treating 10 human GBM lines with 5 µM MT-125 for 48 hours produces multinucleation in 12-25% of tumor cells (**Fig. 1D, E**) (*p<0.0001 for all lines treated with MT-125, compared to vehicle*, *two tailed t test*). To determine if the effects of MT-125 on nuclear size and number can provide a marker of drug-target engagement *in vivo*, we utilized a genetically engineered mouse model (GEMM) (Lei *et al.,* 2011), created by orthotopically injecting a retroviral vector encoding for a PDGFbb-HA fusion protein and for the cre recombinase into mice with a conditional knockout for a tumor suppressor, such as *Trp53* or *Pten*. This produces a proneural, IDH wild type, MGMT unmethylated GBM with 100% penetrance and lethality (Assanah *et al.,* 2006; Lei *et al.,* 2011). We treated mice with *Trp53*(−/−) tumors with vehicle or 10 mg/kg MT-125 administered by IP injection daily for 14 days, sectioned the brains and stained with H&E (**Fig**. 1F & G). MT-125 treatment induces multinucleation (**Fig. 1G**, *white arrows*) and increases mean nuclear cross-sectional area by 53% (**Fig. 1H**, *p<0.0001*).

We also treated GEMM lines deleted for *Trp53*, *Pten*, or both with vehicle or MT-125 for 120 hours, measured cytotoxicity using an ATP-based cell viability assay (*CellTiter-Glo, Promega*), and fit the dose response curves to the Hill equation (**Fig.1I, Table S2**). We performed these same measurements on seven human GBM cell lines, including three (L0, L1, GBM1A) with tumor initiating cell features (Liu *et al.,* 2013; Galli *et al*., 2004) (**Fig. 1J**). This, together with the brain levels reported in our pharmacokinetic studies above and in **Table S1**, predict that doses in the range of 2 to 10 mg/kg should to be efficacious *in vivo*.

Finally, we treated our *Trp53*(−/−) GEMM with vehicle or 10mg/kg IP of MT-125 daily, starting 7 days after retroviral injection, and continued treatment until morbidity. The resulting Kaplan Meier survival curves (**Fig. 1K**) demonstrate that MT-125 significantly prolongs median survival as a single agent (*p<0.0001, log rank test*). When considered with the toxicology data showing the STD 10 is in the range of 40 to 60 mg/kg (**Fig. S3**), efficacy at doses as low as 2 mg/kg (SC) indicate that MT-125 has an excellent therapeutic index of > 20-fold.

### NMII regulates PDGFR signaling

To see if NMII targeting affects RTK signaling, we treated *Trp53*(−/−) cells with 5 µM MT-125 for 48 hours—an exposure time that does not produce appreciable cell death. As illustrated in **Fig. 2A-C**, MT-125 increases expression of PDGFRα and of pY849 PDGFRα 2-3 fold. Deletion of NMIIA also increases PDGFRα expression 7-fold (**Fig. 2D**). MT-125 or deletion of NMII paralogs also increases the phosphorylation of AKT (S473), mTOR (T2446), and S6 kinase (T389) 2-4-fold (**Fig. 2E-I**). Since mTOR is a master regulator of protein translation (Liu and Sabatini, 2020), we wondered if its activation with MT-125 is responsible for enhancing PDGFRα production. To test this, we treated *Trp53*(−/−)cells with MT-125 ± two inhibitors of mTOR function: the TORC1/TORC2 inhibitor AZD8055 and the TORC1 inhibitor everolimus. While neither drug reduces PDGFRα levels below that of vehicle, both prevent MT-125 from increasing PDGFRα expression (**Fig. 2J**). To determine how mTOR activation might accomplish this, we generated a *Trp53*(−/−) GBM cell line that expresses a RiboTag epitope on the Rpl22 ribosomal subunit and treated this line with vehicle or 5 µM MT-125. Using anti-HA pulldowns, we isolated polysomes and measured total and polysomal mRNA encoding PDGFRα using qRT-PCR. Results (**Fig. 2K**) demonstrate that MT-125 specifically enhances polysomal mRNA for PDGFRα by nearly 50% (*p=0.0034, two tailed t test*).

**Figure 2:**
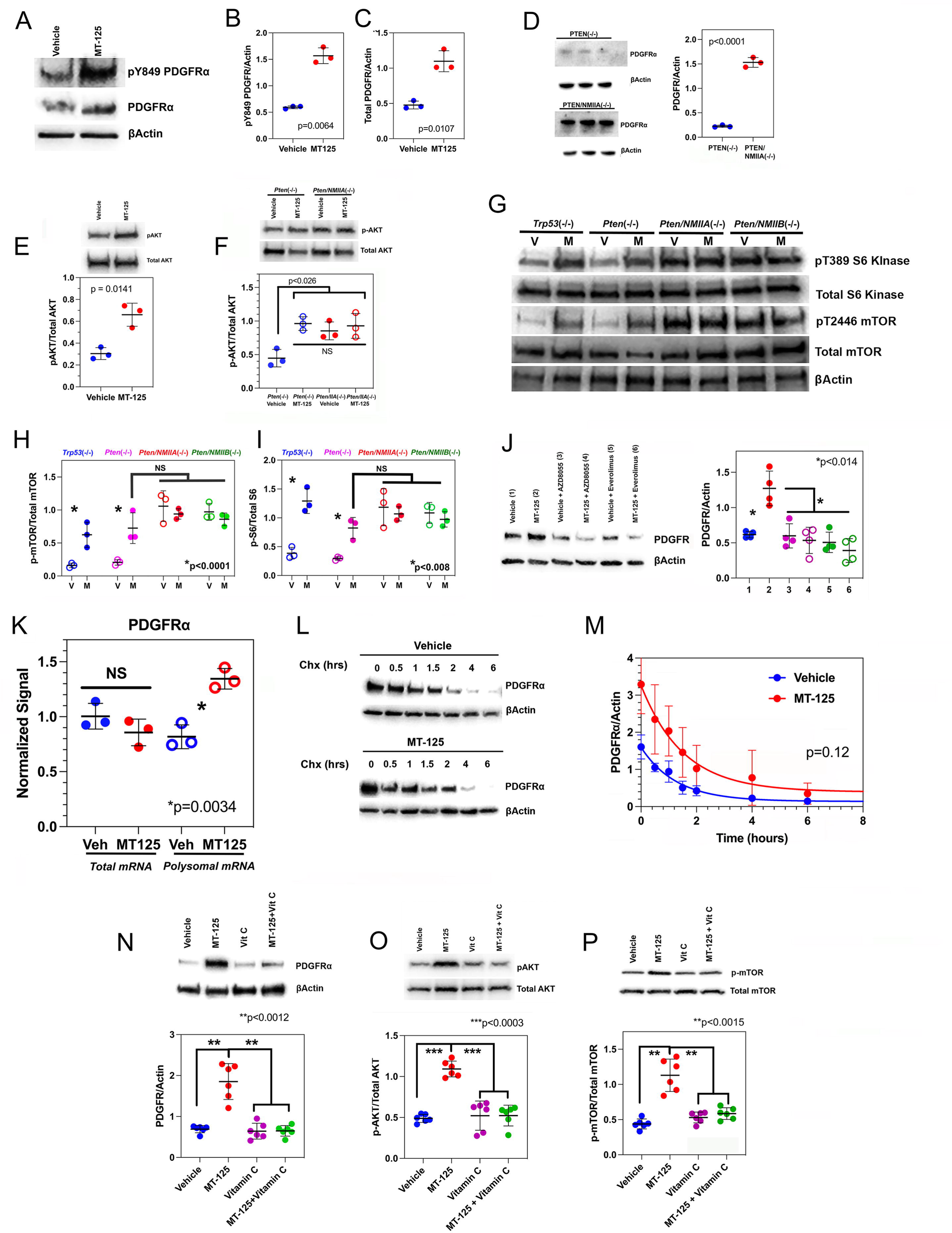
MT-125 regulates signaling along the PDGFR pathway. (**A-C**). Treatment of *Trp53*(−/−) GEMM cells MT-125 increases total and activated PDGFRα 2.5-3-fold (**D**). GBM cells co-deleted for NMIIA and *Pten* increase PDGFRα by >6-fold compared to cells deleted for *Pten*. (**E,F**). MT-125 increases pS473 AKT approximately 2-fold over vehicle in *Trp53*(−/−) (**E**) and *Pten*(−/−) cells (**F**), as does deletion of NMIIA in an *Pten*-deleted background (**F**). (**G**). Levels of S6K, mTOR, pT389 S6K, and pT2446 mTOR were measured in *Trp53*(−/−), *Pten*(−/−), *Pten/NMIIA*(−/−), and *Pten/NMIIB*(−/−) murine GBM cells in the presence of vehicle (V) or MT-125 (M) by Western blot (**G**). (**H,I**). MT-125 increases pS6K and p-mTOR levels in cells deleted for either *Trp53* or *Pten* 2-3-fold over vehicle. The content of pT389 S6K and pT2446 mTOR in *Pten/NMIIA*(−/−) *and Pten/NMIIB*(−/−) GBM cells is increased 3-4 fold compared to *Pten*(−/−) cells. (**J**). The doubling in PDGFRα protein produced by MT-125 (*designated as #2*), compared to vehicle (*designated as #1*), can be blocked by mTOR inhibition with AZD8055 (*designated as #4*) or everolimus (*designated as #6*). (**K**). qRT-PCR of reveals that MT-125 increases polysomal mRNA encoding for PDGFRα ∼50%. (**L**). *Trp53*(−/−)cells were treated with vehicle or MT-125, and PDGFRα was measured by Western *versus* time after adding 10 µM cycloheximide. (**M**). Fitting of data from Fig. 2F for vehicle (*blue*) and MT-125 (*red*) to single exponential decays (*solid curves*) reveals that the rate constants are not significantly different from each other (*p = 0.12)*. (**N-P**). The increase in PDGFRα expression (**N**), and in phosphorylated AKT (**O**) and mTOR (**P**) can be blocked by treatment with 1 mM vitamin C.

The increase in PDGFRα from MT-125 could reflect inhibition of PDGFRα internalization and degradation, due to the role of NMII in receptor mediated endocytosis (Wayt *et al.,* 2021). To test this, we treated *Trp53*(−/−) murine GBM cells with vehicle or MT-125 and measured expression of PDGFRα *versus* time after adding 10 µM cycloheximide to block protein synthesis. Western blot results are depicted in **Fig. 2L** and plots of PDGFRα/βActin intensity ratios *versus* time are depicted in **Fig. 2M**. Fitting data for vehicle (*blue*) and MT-125 (*red*) to single exponential decays (*solid curves*) reveals no statistically significant difference in the slopes of the curves after logarithmic transformation of the data (*p = 0.12*), indicating that MT-125 does not appreciably prolong PDGFRα lifetime. It has also been reported that NMIIA can bind to and inhibit the α subunit of PI3 Kinase (PIK3CA) (Zhang et al., 2021). To test this, we immunoprecipitated lysates from *Trp53*(−/−) GBM cells with an anti-NMIIA antibody, and examined the pulldowns with LC/MS/MS. Our proteomic analysis did not detect any evidence of NMIIA-PIK3CA binding.

Another possible explanation for these findings relates to mitochondrial fission, which utilizes NMII to remove damaged mitochondrial membranes that would otherwise leak toxic reactive oxygen species (ROS) (Korobova *et al.,* 2014; Jun *et al,* 2020). This may explain why inhibiting NMII with a Rho Kinase inhibitor in melanoma was found to increase ROS (Orgaz *et al*., 2020). ROS, in turn, can activate RTK signaling (Corcoran, A and Cotter, TG, 2013; Chiarugi, P and Cirri, P 2003) by oxidizing reactive cysteines in protein tyrosine phosphatases (Lee *et al.,* 1998; Chiarugi *et al.,* 2001). We therefore examined the effect of MT-125 on GBM mitochondria and ROS production, and on the effect of neutralizing ROS on PDGFR signaling. Treatment of *Trp53*(−/−) cells with 5 µM MT-125 for 48 hours increases mean mitochondrial length by >50% (**Fig. S4A** & **B**), doubles the content of ROS-positive cells (**Fig. S4C-E**), and increases the content of mitochondrial ROS by >70% (**Fig. S4F**). We next examined how the anti-oxidant vitamin C affects MT-125 stimulation of PDGFR dependent signaling. We find that the increase in PDGFRα, p-AKT, and p-mTOR induced by MT-125 can be completely reversed by addition of 1 mM vitamin C (**Fig. 2 N-P**).

### NMIIA regulates its own expression and MAPK-dependent signaling

Since we are proposing that NMII regulates activity along the PDGFR-PI3K-AKT-mTOR signaling axis, we wished to see if expression of NMIIA or NMIIB is also regulated by mTOR. We treated both murine and human GBM cell lines with AZD8055 or everolimus, and results are depicted for our *Trp53*(−/−) and *Pten*(−/−) GEMM lines in **Fig. 3A & B** and for two human GBM lines (L0 & L1, described in Galli *et al.,* 2004; Liu *et al.,* 2013) in **Fig. S5 A** & **B**. Both mTOR inhibitors reduce NMIIA approximately 2-fold, while having no significant effect on NMIIB. If NMII suppresses RTK signaling, and if mTOR activation increases NMIIA expression, then by indirectly activating mTOR, MT-125 should also enhance expression of NMIIA. Consistent with this prediction, we find (**Fig. 3C**) that MT-125 increases NMIIA content by ∼60%. We utilized our RiboTag *Trp53*(−/−) GBM line to see if MT-125 alters the distribution of mRNA encoding NMIIA and IIB, analogous to the experiment in **Fig. 2K**. Results (**Fig. 3D & E**) show that while MT-125 increases NMIIA polysomal mRNA by approximately 50%, it has no effect on the corresponding mRNA for NMIIB.

**Figure 3:**
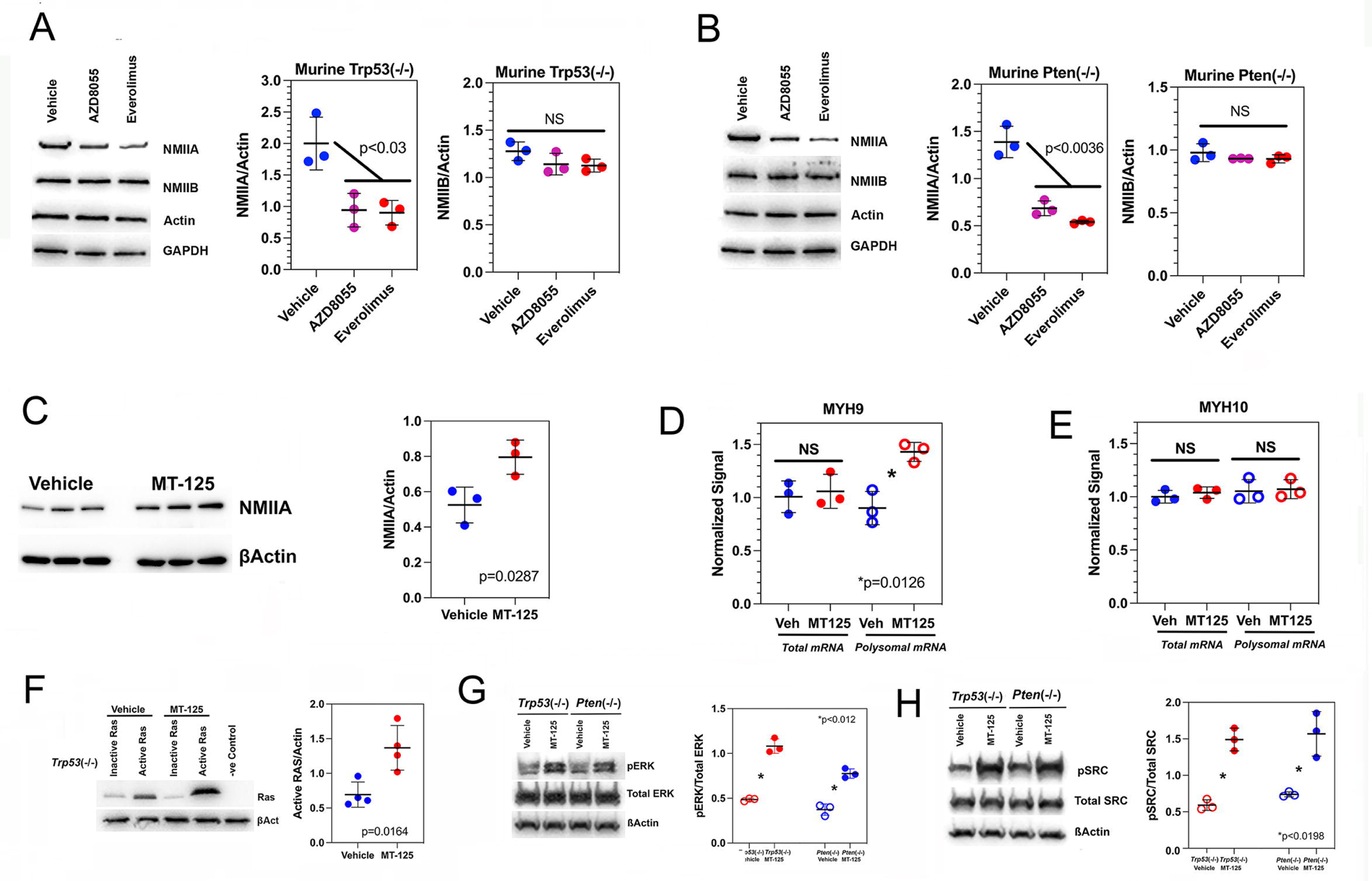
NMIIA expression is regulated by mTOR and regulates the MAPK pathway. (**A,B**). Murine *Trp53*(−/−) (**A**) and *Pten*(−/−) (**B**) cells were treated with vehicle (*blue*), AZD8055 (*magenta*), or everolimus (*red*) for 48 hours and cell lysates were probed for NMIIA and IIB. Both mTOR inhibitors reduce NMIIA protein expression approximately 2-fold, while having no significant effect on NMIIB expression. Very similar results were also seen in two human GBM lines (**Fig. S5**). (**C**). MT-125 produces an ∼60% increase in NMIIA. (**D,E**). MT-125 increases the fraction of translated mRNA encoding for NMIIA (**D**) by ∼50%, while having no significant effect on NMIIB-encoding mRNA (**E**). (**F-H**). Treatment of *Trp53*(−/−) murine GBM cells with 5 µM MT-125 for 48 hours doubles the content of activated RAS (**F**), and similar treatment of both *Trp53*(−/−) and *Pten*(−/−) cells with MT-125 also increases activating phosphorylation of ERK1/2 (**G**) and SRC (**H**) 2-3-fold.

Both RAS and SRC activate the MAPK pathway, and they in turn can be activated through the action of RTKs (Burotto *et al.,* 2014; Lemmon and Schlessinger, 2010). Consequently, we wished to see if MT-125 also stimulates MAPK signaling. Treatment of the *Trp53*(−/−) line with MT-125 doubles the content of activated RAS (**Fig. 3F**), and similar treatment of both *Trp53*(−/−) and *Pten*(−/−) cells with MT-125 also increases phosphorylation of ERK1/2 (**Fig. 3G**) and SRC (**Fig. 3H**) 2-3-fold. We have also examined the effect of MT-125 on oncogenic kinase activation in five low passage human GBM cell lines. Results are summarized in **Fig. S6** and demonstrate that MT-125 enhances activation of PDGFRα, AKT, mTOR, ERK1/2, and SRC to a statistically significanty degree in subsets of these human GBM lines.

#### MT-125 synergizes with oncogenic kinase inhibitors *in vitro*

We have incorporated our results into a model that integrates NMII within two oncogenic signaling pathways, summarized in **Fig. 4A**. We propose that through its role in mitochondrial fission, NMII suppresses formation of ROS, which leads to less PDGFR signaling and less activation of mTOR. TORC1 upregulates expression of PDGFRα (**Fig. 2J,K**), which means that without NMII, there would be a positive feedback loop, with activated mTOR increasing PDGFRα expression, leading to increased mTOR activity, and so on. However, mTOR also increases NMIIA expression, and this arrangement would suppress a PDGFR-mTOR positive feedback loop. By inhibiting NMII, MT-125 would not only increases RTK-PI3K-AKT-mTOR signaling, but also MAPK signaling as well since these two pathways are linked by activating interactions between PDGFRα, SRC, and RAS on the one hand, and between ERK1/2 and mTOR on the other (Burotto *et al.,* 2014; Lemmon and Schlessinger, 2010; Andrae *et al.,* 2008; Carriere et al., 2008). This is consistent with two findings. First, treatment of our *Trp53*(−/−) cells with the PDGFR inhibitor sunitinib reduces not only AKT and mTOR phosphorylation, but phosphorylation of RAS and SRC as well (**Fig. 4B-E**). Second, treatment of these cells with the ERK1/2 inhibitor SCH772984 (**Fig. 4F**) reduces mTOR phosphorylation, even in the presence of MT-125 (**Fig. 4G**).

**Figure 4:**
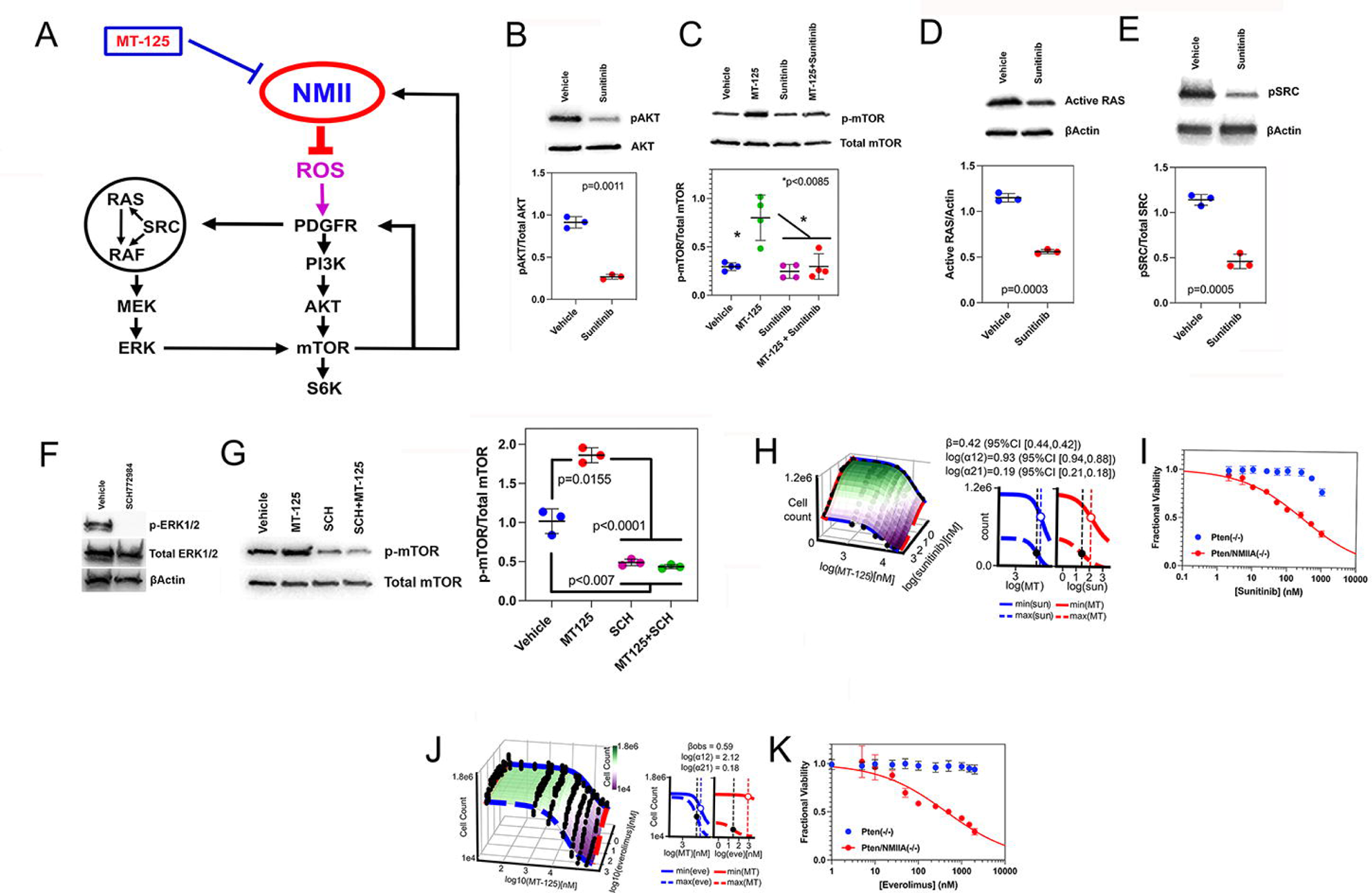
MT-125 synergizes with oncogenic signaling inhibitors *in vitro*. (**A**). Model that incorporates NMII as a negative regulator of RTK-dependent signaling. (**B-E**). Treatment of *Trp53*(−/−) cells with the RTK inhibitor sunitinib reduces pS124 AKT (**B**) and pT2446 mTOR (**C**), activated RAS (**D**) and pY418 SRC (**E**), and in the case of pT2446 mTOR, this effect of sunitinib cannot be overcome by adding MT-125. (**F**). Treatment of *Trp53*(−/−) cells with SCH772984 prevents ERK1/2 phosphorylation at Thr202/Tyr204. (**G**). MT-125-induced increase in pT2446 mTOR can be prevented by SCH772984 (*SHC in figure*). (**H**) (*Left*) Dose-response surface for treatment of Trp53(−/−) GBM cells for 96 hours with combinations of MT-125 and sunitinib. Data for cell count for each pair of drugs (*black dots*) was fit to the MuSyC equation (*surface plot*). The dose-response surface describes the relationship between the drug concentrations (x,y axis) and the cell count (z axis). (*Right*) Projected edges of the surface for the maximum and minimum tested in the combination. Vertical dotted lines mark the EC_50_ for each curve. (**I**). Dose response curves for sunitinib sensitivity for *Pten*(−/−) (*blue*) and *Pten*/NMIIA(−/−) (*red*) cells. Deletion of NMIIA reduces the EC_50_ of sunitinib >10-fold. (**J**). Dose response surface Trp53(−/−) GBM cells treated with combinations of MT-125 and everolimus, analogous to panel **I**. (**K**). Dose response curves for everolimus sensitivity sensitivity for *Pten*(−/−) (*blue*) and *Pten*/NMIIA(−/−) (*red*) cells. Whie NMIIA intact cells are resistant to everolimus, deletion of NMIIA renders them sensitive.

Tumor cells can become dependent on oncogenic pathways that they upregulate, a phenomenon referred to as *oncogene addiction* (Weinstein and Joe, 2008), and this implies that MT-125 might be more effective when combined with inhibitors of RTK-PI3K-AKT-mTOR signaling. We tested this *in vitro* by treating *Trp53*(−/−) GBM cells with combinations of MT-125 + sunitinib and MT-125 + everolimus and measuring the effect on cell count after 96 hours. We analyzed our results with MuSyC, an algorithm that can deconvolve synergy of efficacy from synergy of potency (Wooten *et al.,* 2022; Meyer et al., 2019), and results are depicted as dose response surfaces in the left panels in **Fig. 4I** for MT-125+sunitinib and **Fig. 4K** for MT-125+everolimus. MuSyC fits a dose response surface to drug-combination data to calculate the degree of synergistic efficacy (β) and synergistic potency (log(α12) and log(α21)). MT-125 increases the sunitinib potency by 55% (**Fig. 4I**, log(α21) = 0.19) and everolimus potency by 51% (**Fig. 4J**, log(α21) = 0.18); while MT-125 potency is increased by sunitinib 8.5-fold (**Fig. 4I**, log(α12) = 0.93) and by everolimus 132-fold (**Fig. 4J**, log(α21) = 2.12). Combining MT-125 with sunitinib or everolimus reduces cell count by 42% (β = 0.42) and 59% (β = 0.59), respectively, compared to MT-125, sunitinib, or everolimus alone. Furthermore, genetic deletion of NMIIA also sensitizes *Pten*(−/−) GBM cells to both sunitinib (**Fig. 4I**) and everolimus (**Fig. 4K**). Finally, we screened for synergy between MT-125 and 5 low passage human GBM cell lines, by treating cells with vehicle, MT-125 (5 µM), sunitinib (400 nM), or the combination for 48 hours and measuring cell viability with CellTiter Glo. Each of these lines expresses PDGFRα, although to varying degrees (**Fig. S7 A** & **B**). Combining sunitinib with MT-125 produces a near 2-fold reduction in cell viability in four of the five lines, with a 1.5-fold reduction in the fifth (GBM315), all considerably greater than the effects of single agent MT-125 or sunitinib.

#### MT-125 synergizes with oncogenic kinase inhibitors *in vivo*

We induced *Trp53*-deleted GEMMs by orthotopic retroviral injection as above, and seven days later, randomly assigned mice to receive vehicle (*DMSO*, *daily IP injection*), 10mg/kg MT-125 (*daily IP injection*), 40 mg/kg sunitinib (*oral gavage 5 days/week*), or with the combination of sunitinib and MT-125. Mice were necropsied after 4 weeks of treatment and brain sections obtained at the level of retroviral injection were stained with H&E and an anti-HA antibody to visualize PDGF-secreting tumor cells (**Fig. 5A**). Compared to the vehicle treated sample, whose GBM is highly invasive, the GBM from the MT-125 treated brain is considerably smaller. Sunitinib treatment, as we had previously reported (D’Amico *et al*., 2012), enhances tumor dispersion beyond what is seen with vehicle treatment, with PDGF-positive nests of tumor cells detectable deep within the contralateral hemisphere (*circle in Sunitinib panel*, *enlarged in the right-hand panel*), where they can be seen within pencil fiber white matter tracts (*black arrows in H&E-stained sections*). By contrast, treatment with sunitinib + MT-125 markedly reduces the size of the tumor to an isolated nest of HA-positive cells (*circle in MT-125 + Sunitinib panel*, *enlarged in the right-hand panel*).

**Figure 5:**
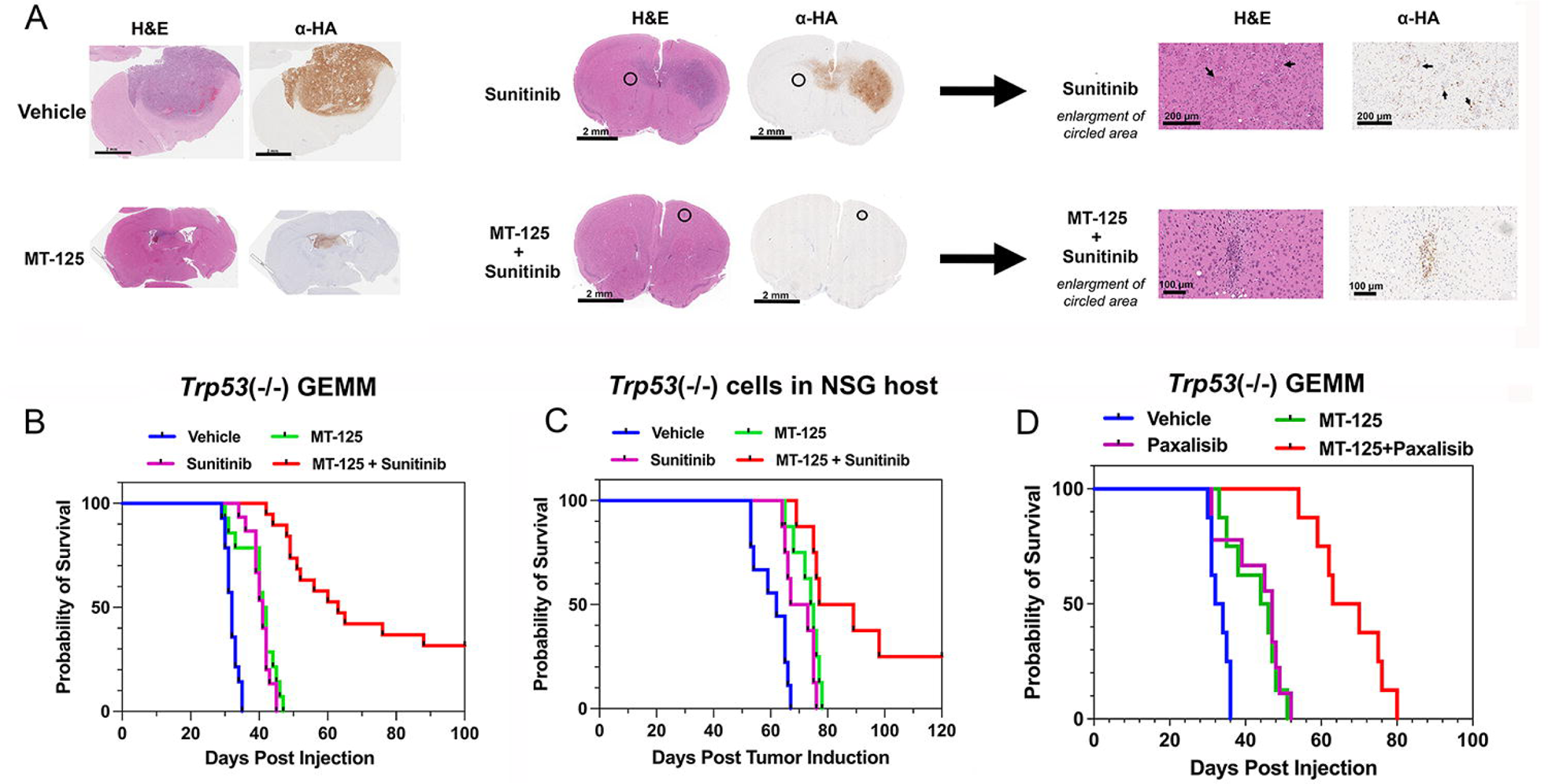
MT-125 synergizes with oncogenic signaling inhibitors *in vivo*. (**A**). *Trp53*(−/−) tumors were treated with vehicle, MT-125, sunitinib, or sunitinib + MT-125. After four weeks of treatment, brain sections were stained with H&E and with an anti-HA antibody to visualize PDGF-secreting tumor cells. Encircled area in sunitinib treated brain shows nests of HA-positive tumor cells in the hemisphere contralateral to the retroviral injection site, with PDGF-positive nests of tumor cells (*black arrows in higher magnification H&E and α-HA stained sections*). Encircled area in the sunitinib + MT-125 treated brain reveals a small focus of tumor cells at the retroviral injection site. (**B**). Kaplan Meier survival curves for mice with orthotopic *Trp53*(−/−) GBMs, treated with vehicle (*blue*), sunitinib (*magenta*), MT-125 (*green*), or sunitinib + MT-125 (*red*). (**C**). NSG mice were orthotopically injected with 50,000 *Trp53*(−/−) cells and after seven days were treated as in (**B**). (**D**). Kaplan Meier survival curves for mice with orthotopic *Trp53*(−/−) GBMs, treated with vehicle (*blue*), paxalisib (*magenta*), MT-125 (*green*), or paxalisib + MT-125 (*red*).

We treated a second cohort of mice with the same drugs and followed them for survival. Monotherapy with either sunitinib (**Fig. 5B**, *magenta*) or MT-125 (**Fig. 5B**, *green*) prolongs median survival over vehicle (**Fig. 5B**, *blue*) by ∼33% (*p<0.0001, log rank test*); with all animals succumbing to tumor. By contrast, combining both drugs (**Fig. 5B**, *red*) prolongs median survival 2-fold over vehicle (*p<0.0001, log rank test*) and produces long term remissions in ∼40% of mice. Dosing with MT-125 + sunitinib stopped at day 85 and no further deaths occurred in this group. In order to estimate how much of the survival benefit from combination therapy depends on lymphoid cells, we established a primary murine *Trp53*-deleted GBM line from the *Trp53*(−/−) GEMM used in **Fig. 5B**, injected these cells orthotopically into NSG mice, and treated them as in **Fig. 5B**. Survival curves are depicted in **Fig. 5C** and show that monotherapy with either sunitinib (**Fig. 5C**, *magenta*) or MT-125 (**Fig. 5C**, *green*) prolongs median survival over vehicle (**Fig. 5C**, *blue*) *(p=0.002 MT-125 vs vehicle, p=0.006, sunitinib vs vehicle, log rank test).* However, the combination of MT-125+sunitinib (**Fig. 5C**, *red*) is still more effective than either alone (*p<0.026, log rank test*), and produces long term remissions in ∼25% of mice.

Our scheme in **Fig. 4A** would also predict that drugs which inhibit oncogenic kinases downstream from PDGFRα should synergize also *in vivo* with MT-125 as well. We tested this in our *Trp53*(−/−) GEMM using paxalisib, a CNS permeant inhibitor of both PI3 Kinase and mTOR (Heffron *et al.,* 2016). As **Fig. 5D** demonstrates, combining paxalisib with MT-125 prolongs median survival to a highly significant degree compared to single agent paxalisib or MT-125 (*p<0.0001, log rank test*).

## DISCUSSION

### NMII paralogs play multiple roles in normal cell physiology that are also relevant to cancer biology

NMII paralogs are essential contributors to cellular function by driving the cytoskeletal dynamics that underlie substrate attachment, cell motility, receptor localization and turnover, and cytokinesis (Straight, *et al.,* 2003). These canonical roles for NMII are also important for tumor cells, as they drive tumor invasion and proliferation, and may explain the the association of NMII upregulation and poorer prognosis in a variety of cancer (Liu *et al.,* 2012; Wang *et al.,* 2021). However, more recent studies have highlighted additional, non-canonical functions of NMII in cell physiology that also are relevant to shaping tumor behavior. These involve a role of NMIIA as a tumor suppressor, through mechanisms that have been reported to include post transcriptional stabilization of *Trp*53 (Coaxum, *et al.,* 2017; Schramek *et al.,* 2014) and inhibition of the PI3 Kinase-dependent signaling (Zheng et al., 2021). In our previous study, we too found that NMIIA functioned as a tumor suppressor in both *Trp53*- and *Pten*-deleted murine GBM models, by inhibiting signaling from several growth and invasion-promoting oncogenic kinases and transcription factors, including SRC, ERK1/2, YAP, and NFκB (Picariello *et al.,* 2019). Since we had previously shown that the *Pten-*deleted GBMs all develop inactivating genetic lesions in *Trp53* or of its downstream effectors (*Lei et al.,* 2011), we conclude that at least in our GBM GEMMs, NMIIA-associated tumor suppression does not require intact *Trp53* function.

### For GBM, a therapeutic strategy targeting NMII needs to include both NMIIA and NMIIB

In our prior study (Picariello, *et al.,* 2019) we also found that while co-deletion of NMIIA and IIB—the two predominant NMII paralogs in GBM—prevented tumorigenesis, deletion of only NMNIIA accelerates tumor growth. We observed that deletion of both NMIIA and IIB is needed to impair cytokinesis in GBM cells, and proposed that the impairment in mitosis from IIA/IIB co-deletion neutralizes the growth promoting effects from deleting just NMIIA. This argues that an NMII-based small molecule therapeutic strategy needs to target both NMIIA and IIB, and this has been the rationale for developing MT-125. We have found that this dual NMIIA/IIB inhibitor is well tolerated with single and repeated dosing regimens at therapeutically relevant doses and achieves therapeutic concentrations with IV, IP, and SC administrations. In addition, it produces nuclear enlargement and multinucleation in orthotopic GBM GEMMs, which could serve as surrogate markers of target engagement in any future clinical investigations of this drug. Furthermore, MT-125 appears to produce little toxicity for up to 85 consecutive days of dosing in mice. We believe that these findings explain how combined NMIIA and IIB inhibition with MT-125 produces a highly statistically significant survival benefit in our aggressive, *Trp53*-deleted GEMM, which validates the potential of MT-125 as a GBM therapeutic.

NMIIA and IIB are widely distributed in vertebrate tissues, which has likely supported an untested assumption that pharmacologic blockade of NMIIA and IIB would be unacceptably toxic. Thus, while our findings that MT-125 is non-toxic and well tolerated refute this assumption, they also raise the question of how normal tissue can tolerate NMII inhibition so much better than GBM does. One possible explanation relates to the findings that NMIIA is upregulated in our GEMMs 2-4-fold (Gill, *et al.,* 2014). If tumor cells are considerably more depencent on NMIIA than are normal cells, then the relatively short half life of MT-125 may still endow this drug with the ability to kill malignant cells while not sufficiently interfering with normal cell function to produce systemic toxicity.

### NMII regulates and is regulated by oncogenic signaling pathways relevant to GBM pathogenesis

Both RTK and MAPK signaling pathways contribute to the aggressiveness of high grade gliomas. We have shown that MT-125 inhibition or genetic deletion of NMIIA activates signaling activity along both pathways, and this has led us to generate a model in which NMII paralogs are integrated within these pathways, functioning as upstream regulators (**Fig. 4A**). In addition to regulating RTK signaling, NMIIA expression is also regulated by this pathway, through the effects of mTOR on the translation of NMIIA-encoding mRNA. We also find that expression of PDGFRα is regulated by mTOR by a similar mechanism. A previous study reported that activation of mTOR suppresses PDGFR-dependent signaling (Zhang *et al.,* 2007) in a set of negative feedback loops. This difference with our findings may reflect corresponding differences in the cell lines used, as that study was performed on cells homozygously deleted for TSC1 and 2, while our murine models express both of these genes (Gill *et al.,* 2014). The relationship between mTOR activity and PDGFRα expression described in **Fig. 4A** would create a reverberating circuit, with PDGFRα-driven activation of mTOR increasing PDGFRα expression, leading to further mTOR activation. However, we propose that NMIIA acts as a brake against runaway RTK pathway activation. Our model would also predict that this homeostatic mechanism breaks down with MT-125 inhibition of NMII, so the consequent sustained activation of both PDGFR and MAPK signaling would lead to increased dependency on these growth promoting pathways, and enhanced sensitivity to drugs that target RTK pathway inhibitors. This conclusion is consistent with our earlier findings (Picariello et al., 2019) that genetic deletion of NMIIA in GBM models enhances ERK1/2 activity, and induces sensitivity to an ERK1/2 inhibitor which in NMIIA intact tumors has little effect.

NMII paralogs play important roles in driving receptor mediated endocytosis (Wayt *et al.,* 2021), a process that can limit the activity of an RTK by helping to traffic it to lysosomes for degradation. However, our data is not consistent with MT-125 affecting the lifetime of total PDGFRα. Likewise, while NMIIA-mediated binding to and inhibition of PIK3CA could have also provided a plausible mechanism for explaining the effects of MT-125 (Zheng et al. 2021), we cannot detect interactions between these two proteins in our GBM models. Instead, our data support an NMII-dependent mechanism that involves the effects of ROS on oncogenic kinase signaling. During mitochondrial fission, NMII paralogs form a contractile ring around damaged mitochondrial segments, analogous to what occurs in cytokinesis, and extrude these segments for degradation in the lysosomes in a process called mitophagy (Onishi *et al.,* 2021). Mitochondrial fission is a quality control mechanism that is designed to reduce leakage of ROS into the cytoplasm and nucleus from damaged mitochondria (Korobova *et al.,* 2014). Consistent with this mechanism, we find that inhibiting NMII with MT-125 lengthens mitochondria and increases cytoplasmic and mitochondrial ROS. However, ROS can also function as a signaling molecule (Corcoran, A & Cotter, TG, 2013; Chiarugi, P & Cirri, P 2003) and in particular, can activate RTKs, possibly by inactivating protein tyrosine phosphatases (Chiarugi, P and Cirri, P, 2003; Chiarugi et al., 2001). The concept that a biologically reactive gas molecule can regulate signalling is hardely novel, as nitric oxide is a well documented example of a gas that modulates signaling, at least in part through its ability to modify amino acid side chains through both nitrosylation and tyrosine nitration (Moriguchi *et al.,* 1992; Handy and Loscalzo, 2006).

### MT-125 represents a first-in-class small molecule inhibitor that has significant potential for the treatment of GBM

The prognosis for patients afflicted with GBM has remained dismal in spite of decades of clinical investigative efforts (Bai *et al.,* 2011; Mellinghoff, *et al.,* 2012), which speaks to the need to find novel targets with inhibitors that are well tolerated, are brain permeant and retained in the CNS, and which are not only effective on their own but which synergize with other CNS permeant, FDA approved inhibitors as well. We propose that MT-125 meets these criteria. This drug is well tolerated in a variety of pre-clinical animal models, with minimal hematologic or metabolic toxicities, is active against both murine and human GBM cell lines, and prolongs survival as a single agent in a murine GBM GEMM. Furthermore, consistent with our earlier studies (Picariello et al., 2019), we also find that MT-125 synergizes with an FDA approved, CNS permeant RTK inhibitor (sunitinib), with the combination producing long term survival in a highly aggressive GBM model. Our results provide powerful rationale for developing NMII targeting strategies to treat cancer and demonstrate that MT-125 has strong clinical potential for the treatment of GBM.

## ACKNOWLEDGEMENTS

We wish to thank Drs. Alfredo Quinones-Hinojosa and Hugo Guerrero-Cazares (Mayo Clinic Florida) for their gift of the 1A, 108, 120, 612, 166, 315, 509, and 612 human primary GBM cell lines, Dr. Dolores Hambardzumyan (Mount Sinai School of Medicine) for her gift of the MES1861 murine GBM cell line, Dr. Justin Lathia (Cleveland Clinic Foundation) for his gift of the L0 and L1 cell lines, and Susan Khan for technical assistance. S.S.R. is supported by NIH grants NS073610, NS118513, NS119714, and CA210910 and by a Translational Adult Glioma Award from the Ben and Catherine Ivy Foundation. P.C. is supported by NS118513, CA251313, and CA250481. The work was further supported by R01DA049544 to CAM and UH3NS096833 to CAM, PRG and TMK. CTM was supported by the K99/R00 Pathway to Independence Award from NIAID (1K99AI175656).

## AUTHOR CONTRIBUTIONS

Conception and design: RK, LR, EJY, NZ, LL, AD, PC, MDC, GR, PRG, TMK, CAM, and SSR Development of methodology: RK, LR, NZ, AD, CTM, PC, PRG, TMK, CAM, and SSR Acquisition of data: RK, LR, EJY, NZ, LL, AD, AH, MDC Analysis and interpretation of data: RK, LR, EJY, NZ, AD, CTM, PC, GR, PRG, TMK, CAM, and SSR

## DECLARATION OF INTERESTS

Erica Young, Patrick Griffin, Theodore Kamenecka, Courtney Miller and Steven Rosenfeld have an equity interest in Myosin Therapeutics. Christian T. Meyer is a co-founder of Duet Biosystems. No other authors have competing interests to declare.

## METHODS

### Materials availability

Genetically engineered mouse models and cell lines will be distributed after completion of the relevant Materials Transfer Agreements with the Mayo Clinic.

### Data and code availability

The datasets and code used in this study are publicly available through the publications referenced in this study and in the following sections.

### Kinetic aqueous solubility

Kinetic aqueous solubility of MT-125 was determined as reported previously (PMID: 34558887).

### Protein Isolation and Purification

Bovine CMII was purchased from Cytoskeleton, Inc (cat. MY03). Human NMIIA was isolated from platelets and thiophosphorylated as reported previously (PMID: 34558887). F-actin was prepared from rabbit Rabbit muscle acetone powder (cat. 41995-2, Pel Freez Biologicals) according to the protocol of Pardee and Spudich (PMID: 7098993).

### NADH-coupled ATPase assays

The potency of MT-125 against NMIIA and CMII was assessed in a fluorescence-based nicotinamide adenine dinucleotide (NADH)-coupled ATPase assay. A detailed protocol has been reported elsewhere (PMID: 31475972). The assay concentration of CMII, NMIIA, and actin was 300 nM, 400 nM, and 10 µM, respectively.

### Cytokinesis Assay

The potency of MT-125 on NMIIB was determined in a COS7 cell-based cytokinesis assay as described previously (PMID: 33204739).

### Animals

Adult, 8-10-week-old male C57BL/6 mice (25-30 g, Jackson Laboratory) were housed under a 12:12 light/dark cycle, with food and water ad libitum. Adult male and female 225-300 g rats (Charles River) at the start of the experiment were handled and housed under a 12:12 reverse light/dark cycle. All procedures at UF Scripps were performed in accordance with the UF Scripps Research Institutional Animal Care and Use Committee.

All mouse procedures at the Mayo Clinic were performed in compliance with the Mayo Clinic Institutional Animal Care and Use Committee guidelines. Homozygous floxed *Trp53* mice (Stock #008462) and NOD-SCID mice (Stock #005557) were obtained from Jackson Laboratory. Studies were performed on equal numbers of male and female mice. Animal genotypes were regularly verified via tail snip (TransnetYX, Cordova, TN).

### Drugs

Blebbistatin (Tocris, Minneapolis, MN), MT-125 (Myosin, Jupiter, FL). The vehicle for MT-125 was 30% HPBCD, 50 mM Acetate Buffer pH 5.

### Pharmacokinetics

Blood was collected from rats or mice at various time points post infusion (see Table S1) into lithium heparin coated tubes (cat. 07 6101, Ram Scientific), and stored on wet ice. Blood samples were later centrifuged for 3 min at 2,655 g to separate plasma from red blood cells. Plasma was collected into a fresh tube and stored at −80 °C. In addition to blood samples, temporal lobe brain tissue was collected from each mouse. Each sample was flash frozen with 2-methylbutane and stored at −80 °C. Compound levels were quantified in brain and plasma by mass spectrometry using an ABSciex 5500 mass spectrometer using multiple reaction monitoring. Brain samples were homogenized in water and then immediately treated with 5-times (v:v) acetonitrile to extract the compound and precipitate cellular protein. Plasma samples were directly treated with acetonitrile. Samples were filtered through a 0.45 µm filter plate prior to injection onto the LC-MS/MS.

### Glioma cell line isolation from mouse GBM tumor and culture

The protocol for isolation of tumor cells from *Trp53(−/−)* and *Pten(−/−,), Trp53/Pten(−/−). Pten/NMIIA(−/−), and Pten/NMIIB(−/−)* murine tumors has been described (Lei *et al*. 2011). The human L0 and L1 primary GBM cell lines were cultured and maintained in DMEM+F12 media with 1% N2 supplement (Gibco), 20ng/ml of hEGF (Sigma-Aldrich) and 20ng/ml of hFGF (R&D systems). Human GBM cells 1A, 108,120, 612, 509, 593,166, and 315 primary lines were cultured and maintained in DMEM+F12 media with 1% NeuroPlex supplement (Gemini), 20ng/ml of EGF and 20ng/ml of FGF. Murine mesenchymal (MES) glioblastoma cells (MES1861 and MES4622) which lack expression of *Nf1* and *Trp53* were maintained in DMEM media with 10% FBS as previously described (Reilly *et al.,* 2000; Gursel *et al.,* 2011).

### Dose response curves/cell viability assays

5000 cells/well were plated in 96-well plates were treated for 96 hours with various doses of MT-125, sunitinib, everolimus, or combinations of MT-125 and sunitinib or everolimus. Cell counts per well were measured using CellTiter Glo (Promega, cat# G9242).

### Transwell in vitro invasion assays

Fluoroblok Transwell inserts added with 125,000 cells for each insert. 10% FBS was used as a chemoattractant. Cells were incubated for 15 hours at 37°C, inserts were washed with PBS, fixed in 4% PFA for 15 min and washed twice with PBS before staining with DAPI. Images were captured and analyzed for nuclear count using Cytation 5.

### Retrovirus production and intracerebral injections

PDGF-IRES-cre retrovirus was generated and injected intracranially into mice floxed for TP53 according to methods described previously (Lei *et al.,* 2011; Kenchappa *et al.,* 2020). For the pharmacologic studies mice were treated seven days after retroviral injection with vehicle, sunitinib (40 *mg/kg by oral gavage, 5 days per week*), MT125 (*10 mg/kg by intraperitoneal injection everyday*), MT125 + sunitinib (MT125*: 10 mg/kg by intraperitoneal injection for everyday; sunitinib:* 40 mg/kg *by oral gavage, 5 days per week*), paxalisib (8 *mg/kg by oral gavage, 5 days per week*) or MT125 + sunitinib (MT125*: 10 mg/kg by intraperitoneal injection for everyday; paxalisib: 8* mg/kg *by oral gavage, 5 days per week*) with 7-15 mice per treatment group. Treatment continued until tumor morbidity. For studies using the *Trp53*(−/−) GBM model, NOD/SCID mice were orthotopically injected with 100,000 *Trp53*(−/−) GBM cells and treatment with vehicle, MT-125, sunitinib, or MT-125 + sunitinib began seven days later. Treatment was continued until tumor morbidity.

### Western blots

Cells were scraped and incubated in lysis buffer (50 mM Tris HCl at pH 7.40, 150 mM NaCl, 1 mM EDTA, 1.0% Nonidet P-40, and a mixture of protease and phosphatase inhibitors), on ice from 30 minutes. Debris was removed by centrifugation for 10 minutes at high speed at 4°C, and cleared lysates were run on SDS/PAGE and transferred to polyvinylidene difluoride membranes. Membranes were blocked in 5% non-fat dry milk in TBS + 0.1% Tween 20 for 1 hour at room temperature, incubated with primary antibody in blocking solution for overnight at 4°C, followed by secondary antibody for 1 hour at room temperature, and developed using an enhanced chemiluminescence solution.

### RAS activity assay

RAS activity was assessed by affinity pull-down of endogenous levels of GTP-bound Active-Ras as described by Active Ras detection kit (Cell Signaling Technology, cat# 8821).

### Brain histological analysis

Brains from 4% paraformaldehyde-perfused, GBM-bearing mice were paraffin-embedded as described (Kenchappa *et al.,* 2020). IHC was performed on 5 mm sections using the Discovery ULTRA automated stainer (Ventana Medical Systems). Antigen retrieval was performed using a Tris/borate/EDTA buffer (Discovery CC1), pH 8.0-8.5, for 60 minutes at 95°C. Slides were incubated with anti-HA for 2 hours at room temperature. The antibodies were visualized using biotinylated goat anti-rabbit and rabbit anti-rat secondary and counterstained with hematoxylin and eosin (H&E). Images were captured and analyzed using a ScanScope scanner and ImageScope software (Aperio Technologies).

### Drug synergy

Synergy was calculated using the MuSyC algorithm as previously described (Wooten et al., 2021). MuSyC quantifies two types of drug synergy, synergistic potency and synergistic efficacy, both relating to geometric transformations of the dose response surface, which are analogous to the transformations in the 1D Hill equation for potency (horizontal shift in the EC50) and efficacy (vertical shift in Emax). Synergy was calculated by fitting a dose-response surface relating the drug’s effect (cell count at 96 hours) to the concentrations of drug 1 and drug 2.

### NMIIA pulldown

The p53(−/−) cells underwent a 24-hour exposure to either the Vehicle or SR561. Subsequently, cellular proteins were extracted through the utilization of RIPA lysis buffer, supplemented with protease and phosphatase inhibitors. Specific antibodies directed against NMIIA or IgG were incubated overnight at 4°C with 1000μg of total cellular lysates. Following this, samples underwent a 1-hour incubation at room temperature with Protein A magnetic beads, succeeded by meticulous washing to eliminate non-specific interactions. The magnetic beads were then resuspended in 500μL of ultrapure water. A western blot analysis for NMIIA was executed using 50μL of the final bead suspension in water to validate the immunoprecipitation outcomes. The processed samples were subsequently forwarded to the Proteomics core for a comprehensive analytical assessment.

### Polysomal pulldown and RT-qPCR

The polysomal pulldown assay was conducted utilizing BC7 cells exhibiting a specific genotype characterized by PDGFB overexpression, p53 deletion, and RiboTag labeling (REF of cell line from Peter Canoll lab). All reagents employed were RNAse-free. A total of 5 million cells underwent pre-incubation with 50 µg/mL Cycloheximide for 5 minutes to mitigate polysomal degradation during the harvest process. Experimental conditions encompassed treatment with Vehicle and SR561 for 48 hours. Cell lysis was carried out using 1 mL of Polysome Lysis Buffer (PLB) supplemented with RNase inhibitor and DNase. An aliquot of 100 µL of the lysate served as the input and underwent RNA purification with the RNeasy Mini kit. Concurrently, 450 µL of the lysate was subjected to overnight incubation with pre-washed anti-HA magnetic beads at 4°C with rotation. Beads were subsequently washed four times with PLB, followed by the addition of 450 µL RLT buffer. The beads were then separated using a magnet, and the supernatant underwent RNA purification with the RNeasy Mini kit. The quantified RNA (500 ng) was utilized for cDNA production utilizing iScript cDNA synthesis kit. Subsequently, qPCR was performed to determine the relative mRNA expression levels of PDGFRa, NMIIA (Myh9), and NMIIB (Myh10) using specific primers and SYBR Green. PRLP0 expression served as the housekeeping gene. Three independent experiments were conducted, and input samples were included as controls.

### Ferroptosis quantification

Ferroptosis was evaluated using confocal microscopy and flow cytometry methodologies, employing MitoSOX and MitoCLOX reagents in L1 and p53 (−/−) cells after a 48-hour treatment with either the Vehicle or SR561. In the confocal microscopy protocol, cells were cultivated in glass-bottom chambers slides. Following the treatment, the culture media were aspirated, and a complete media containing MitoSOX or MitoCLOX (diluted at ratios of 1:2000 and 1:5000, respectively) was applied for 30 minutes. Hoechst 33342 was then introduced for 5 minutes exclusively for MitoSOX assays. After two washes with PBS, complete media were reinstated, and cellular imaging was executed utilizing the Zeiss LSM 880 microscope with a 20x objective, capturing 10-15 images in total per experimental condition. Quantitative fluorescence analysis was carried out using FIJI software. MitoSOX fluorescence levels were normalized with reference to Hoechst 33342 staining. For MitoCLOX quantification, fluorescence measurements at 520 nm were recorded to assess an increase, while the initial fluorescence at 590 nm was examined for a decrease. In the context of flow cytometry, cellular populations were harvested post-treatment with Vehicle or SR561. Subsequently, 1 million cells were exposed to MitoSOX or MitoCLOX for 30 minutes. Sample data were collected using a Cytoflex flow cytometer to record the percentage of staining cells and mean fluorescence intensity. Data files were subjected to analysis using FlowJo v10.9. The entire procedure was systematically replicated in three independent experiments.

### Expression levels of PDGFRα

The expression of PDGFRa was evaluated at transcriptional and translational levels using RT-qPCR and Western blot techniques, respectively. PTEN (−/−) and PTEN/NMIIA (−/−) cells cultured in 6well plates at 70% confluency were utilized for the study, and three independent experiments were conducted to ensure the robustness of the findings. RNA extraction was performed using the TRIzol plus RNA purification kit for the RT-qPCR analysis. The quality and quantity of RNA samples were determined with Nanodrop, followed by cDNA synthesis using the iScript kit with 500 ng of total RNA. Primers targeting PDGFRa and PRLP0 mRNAs were employed for qPCR, and the QuantStudio3 real-time machine, utilizing SYBR Green, was used for qPCR amplification. The real-time PCR data files were analyzed using QuantStudio and Analysis software. In the case of Western blot analysis, protein lysis was carried out using RIPA buffer containing proteinase and phosphatase inhibitors. The total protein concentration was determined using the BCA assay, and 100 µg of total protein was loaded for each sample. An anti-PDGFRα antibody was used for Western blotting, and the membrane was developed using ECL, with images captured using a bioimager. The intensity of the bands in the Western blot was quantified using FIJI software, with β-actin serving as the normalization reference.

### Integrin β1 immunofluorescence

The active levels of Integrin β1 were determined through immunofluorescence analysis in p53 (−/−) cells subjected to a 72-hour treatment with either Vehicle or SR561. The cells were seeded in glass bottom chamber slides, followed by PBS washing post-treatment. Subsequently, fixation was performed using 4% paraformaldehyde for 20 minutes, succeeded by a 1-hour blocking step with 10% normal goat serum. Total integrin β1 and active integrin β1 primary antibodies were then added, and the samples were incubated overnight at 4°C. The following day, after the removal of primary antibodies and washing, a secondary antibody (Alexa Fluor 647) was added and incubated for 1 hour. Further washing was conducted, and a solution of Rhodamine Phalloidin with DAPI was applied for 10 minutes. After a subsequent PBS wash, the samples were imaged using a Zeiss LSM 880 Confocal microscope with AiryScan technology, employing a 45x oil objective. Fluorescence intensity was quantified utilizing FIJI software, with the fluorescence at 647 (indicative of active Integrin β1) normalized against the fluorescence at 488 (total Integrin β1). This entire process was replicated in three independent experiments.

### Data analysis

For *in vitro* studies, a two tailed t test or one-way ANOVA was used to calculate p values with statistical significance at p<0.05. For survival studies, statistical significance was determined using a log rank test, and significance was set at p<0.05.

## Notes

### Competing Interest Statement

The authors have declared no competing interest.

