## Supplement for "MT-125 Inhibits Non-Muscle Myosin IIA and IIB, Synergizes with Oncogenic Kinase Inhibitors, and Prolongs Survival in Glioblastoma"

### SUPPLEMENTARY TABLES

#### Table S1. In Vivo Pharmacokinetic Analysis of MT-125

| | Plasma | | | | Brain | | | |
|---|---|---|---|---|---|---|---|---|
| Dose (mg/kg) | $C_{max}$ (μM) | $T_{max}$ (hr) | $T_{½}$ (hr) | $AUC_{last}$ (μM.hr) | $C_{max}$ (μM) | $T_{max}$ (hr) | $T_{½}$ (hr) | $AUC_{last}$ (μM.hr) |
| **5** | 3.8 ± 0.2 | 0.2 ± 0.0 | 12.4 ± 2.5 | 4.2 ± 0.2 | 9.1 ± 1.0 | 0.3 ± 0.1 | 10.5 ± 1.4 | 5.4 ± 0.8 |
| **10** | 9.0 ± 0.2 | 0.2 ± 0.0 | 17.6 ± 4.9 | 12.5 ± 0.2 | 7.5 ± 1.0 | 0.2 ± 0.1 | 15.7 ± 2.8 | 7.5 ± 0.1 |

Values represented are means ± SEM

**Table S2: EC50 and Hill Coefficients for MT-125 Cytotoxicity in GBM Cell Lines**

| Cell Line | Source | $EC_{50}$ (nM) | Hill Coefficient |
|---|---|---|---|
| *1A* | Human | 3315 ± 315 | 1.00 |
| *L0* | Human | 3523 ± 487 | 1.02 |
| *L1* | Human | 3411 ± 208 | 0.80 |
| *612* | Human | 3749 ± 301 | 0.94 |
| *120* | Human | 4236 ± 320 | 0.78 |
| *166* | Human | 4554 ± 290 | 0.94 |
| *315* | Human | 3644 ± 287 | 0.85 |
| *p53(-/-)* | Murine | 3924 ± 235 | 1.43 |
| *PTEN(-/-)* | Murine | 7232 ± 255 | 1.52 |
| *p53/PTEN(-/-)* | Murine | 3828 ± 201 | 1.40 |

### SUPPLEMENTARY FIGURES

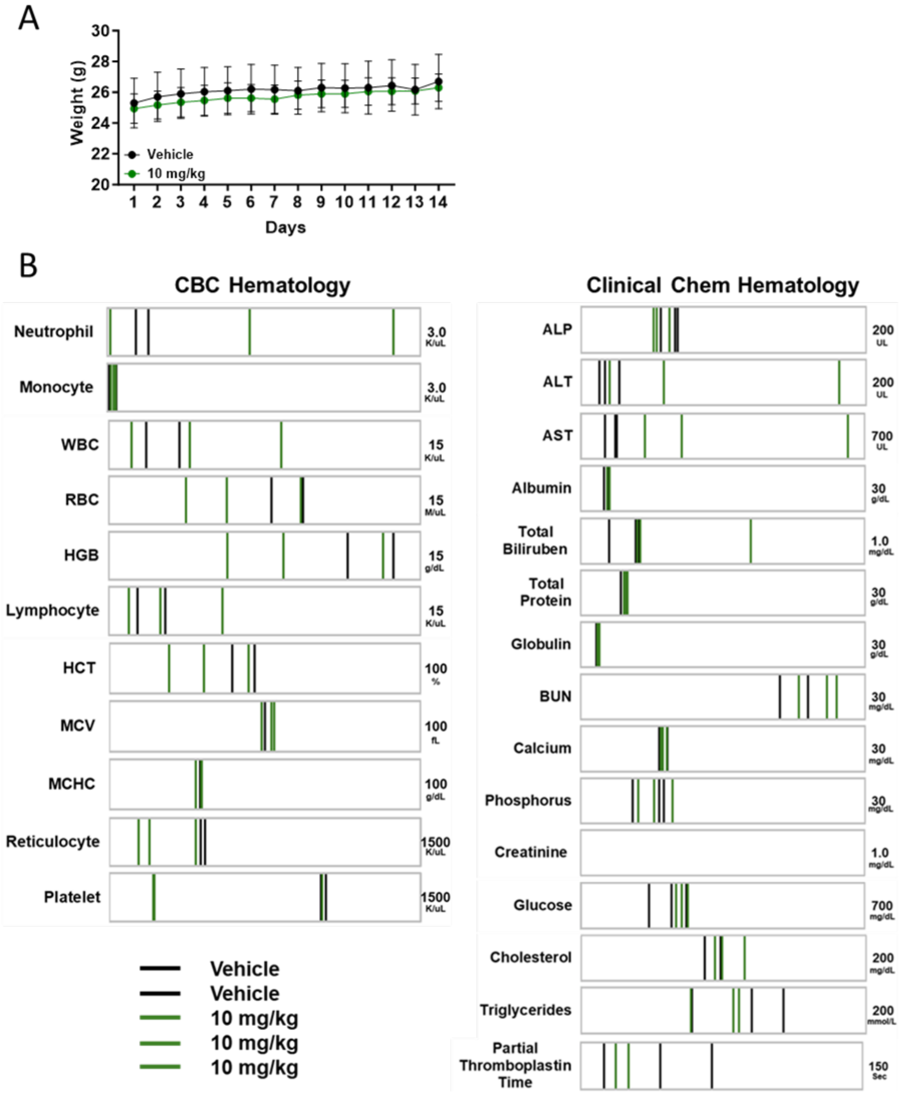

**Figure S1. Repeated MT-125 dosing results in no adverse health effects.** Mice treated daily with 10 mg/kg MT-125 SC for 14 days had no significant (A) change in weight or (B) clinical chemistry and hematology compared to vehicle treated animals.

A

| Group | Weight | | Averages | |
|---|---|---|---|---|
| | Day 1 | Day 2 | Day 1 | Day 2 |
| 60 mpk | 348 | 346 | 350.33 ± 6.74 | 350.0 ± 7.21 |
| | 340 | 340 | | |
| | 363 | 364 | | |
| 70 mpk | 352 | 352 | 366.33 ± 12.86 | 370.67 ± 14.85 |
| | 392 | 400 | | |
| | 355 | 360 | | |
| 90 mpk | 328 | 324 | 363.33 ± 17.90 | 364.0 ± 20.01 |
| | 376 | 383 | | |
| | 386 | 385 | | |

B

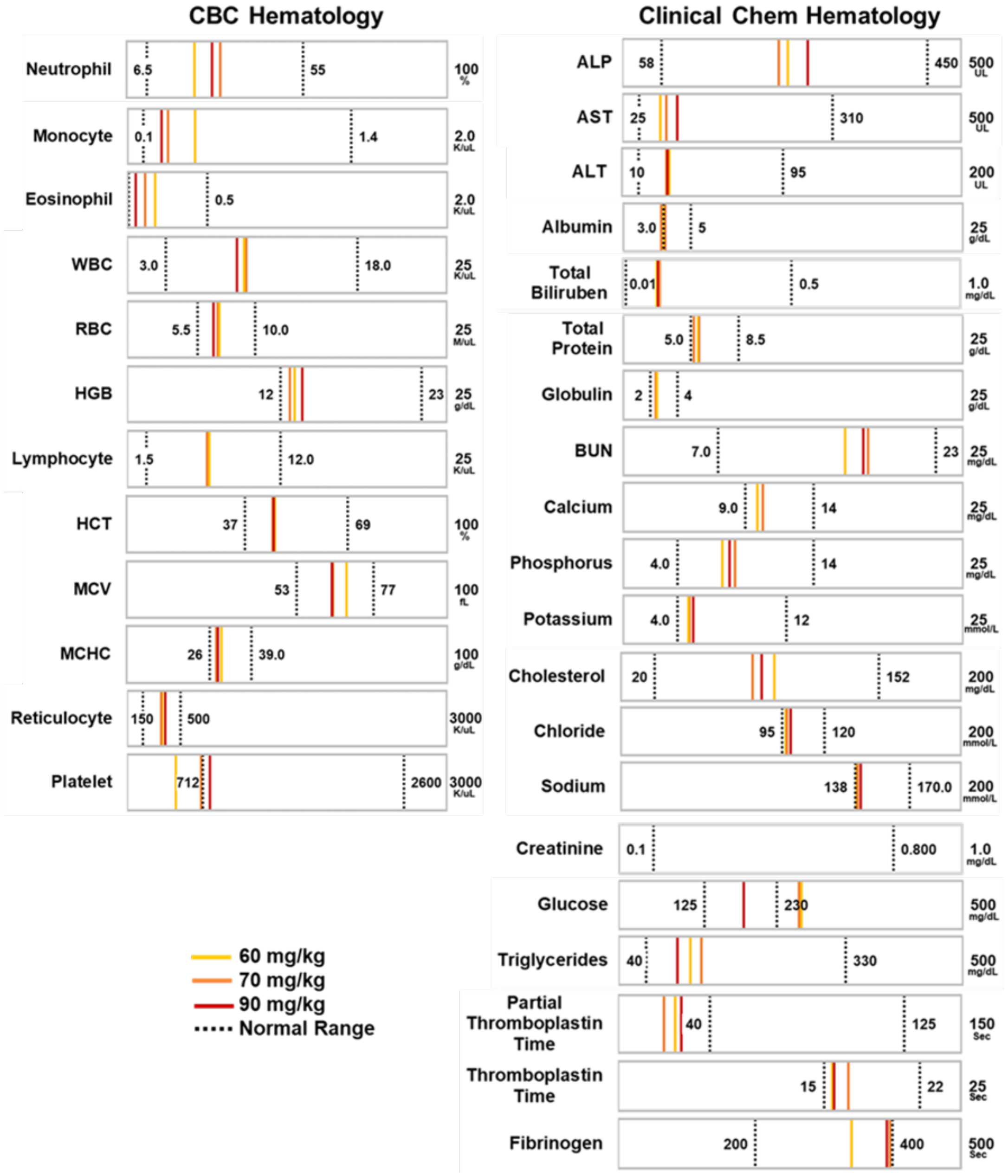

**Figure S2. Single doses as high as 90 mg/kg were well tolerated by rats.** Animals were injected with 60, 70 or 90 mg/kg MT-125 SC and 24 hrs later showed no significant change in (A) weight or (B) clinical chemistry and hematology.

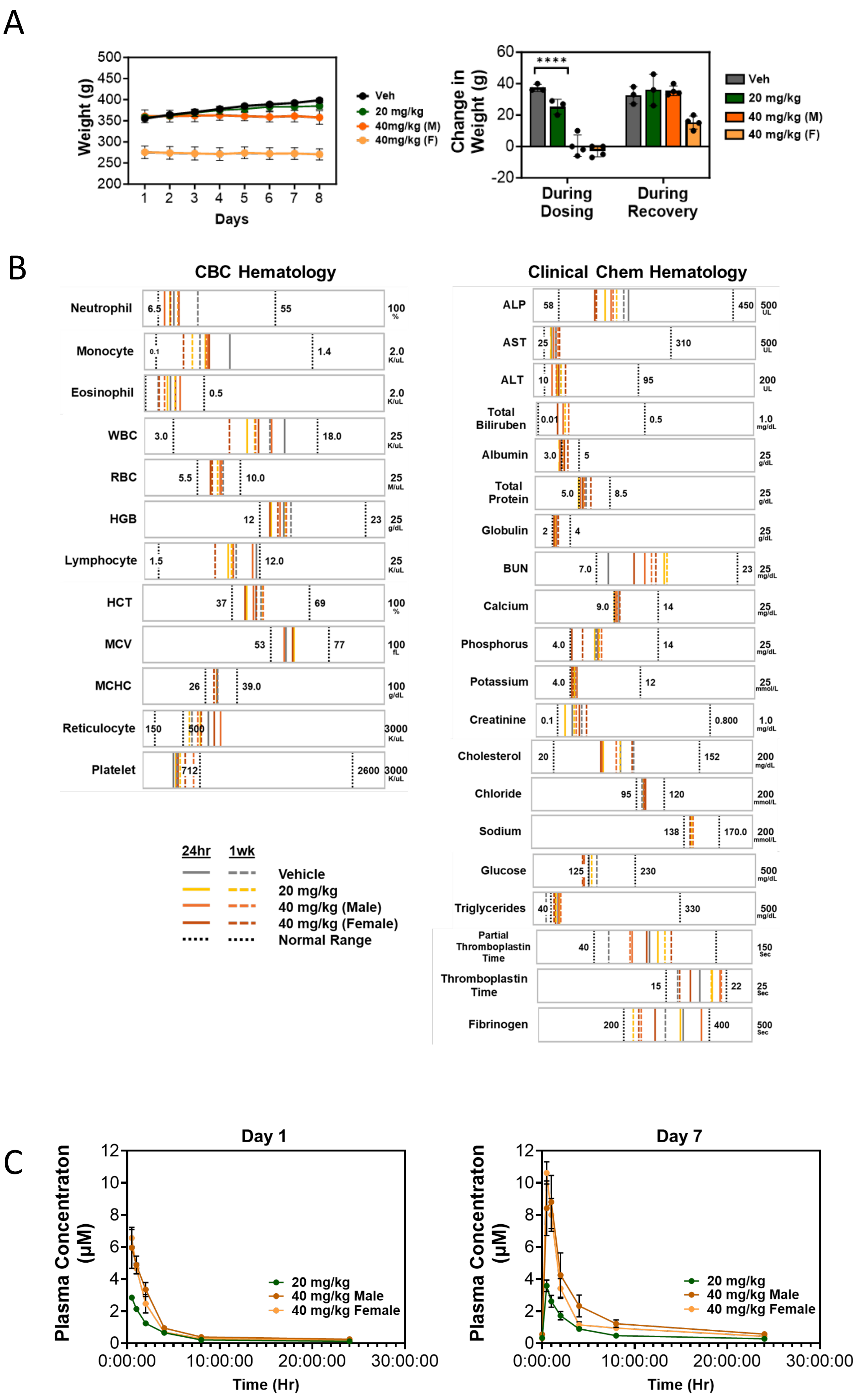

A
Weight (g)
Days
Veh
20 mg/kg
40mg/kg (M)
40mg/kg (F)
Change in Weight (g)
****
During Dosing
During Recovery
Veh
20 mg/kg
40 mg/kg (M)
40 mg/kg (F)
B
CBC Hematology
Neutrophil
Monocyte
Eosinophil
WBC
RBC
HGB
Lymphocyte
HCT
MCV
MCHC
Reticulocyte
Platelet
Clinical Chem Hematology
ALP
AST
ALT
Total Biliruben
Albumin
Total Protein
Globulin
BUN
Calcium
Phosphorus
Potassium
Creatinine
Cholesterol
Chloride
Sodium
Glucose
Triglycerides
Partial Thromboplastin Time
Thromboplastin Time
Fibrinogen
24hr
1wk
Vehicle
20 mg/kg
40 mg/kg (Male)
40 mg/kg (Female)
Normal Range
C
Day 1
Day 7
Plasma Concentraton (µM)
Time (Hr)
20 mg/kg
40 mg/kg Male
40 mg/kg Female

**Figure S3. *In vivo* toxicology testing of MT-125.** Rats received daily doses of vehicle or MT-125 (20 or 40 mg/kg, SC) for seven days. (**A**) Weight was monitored daily during the administration period and following the 7-day recovery period. Dosing 40 mg/kg, but not 20 mg/kg, attenuated age-related weight gain during dosing but resumed during recovery period (2-way rmANOVA $F_{(21, 70)} = 19.6$ $p < 0.0001$; Day 1 vs Day 8 Veh: $p < 0.01$, 20 mg/kg: $p < 0.01$; Change in weight 2-way ANOVA $F_{(3,10)} = 20.2$, $p < 0.0001$). (**B**) Clinical chemistry and hematology samples were collected on Day 8 (dosing period) and 14 (recovery period). (**C**) Toxicokinetic timecourse samples were collected on Days 1 and 7.

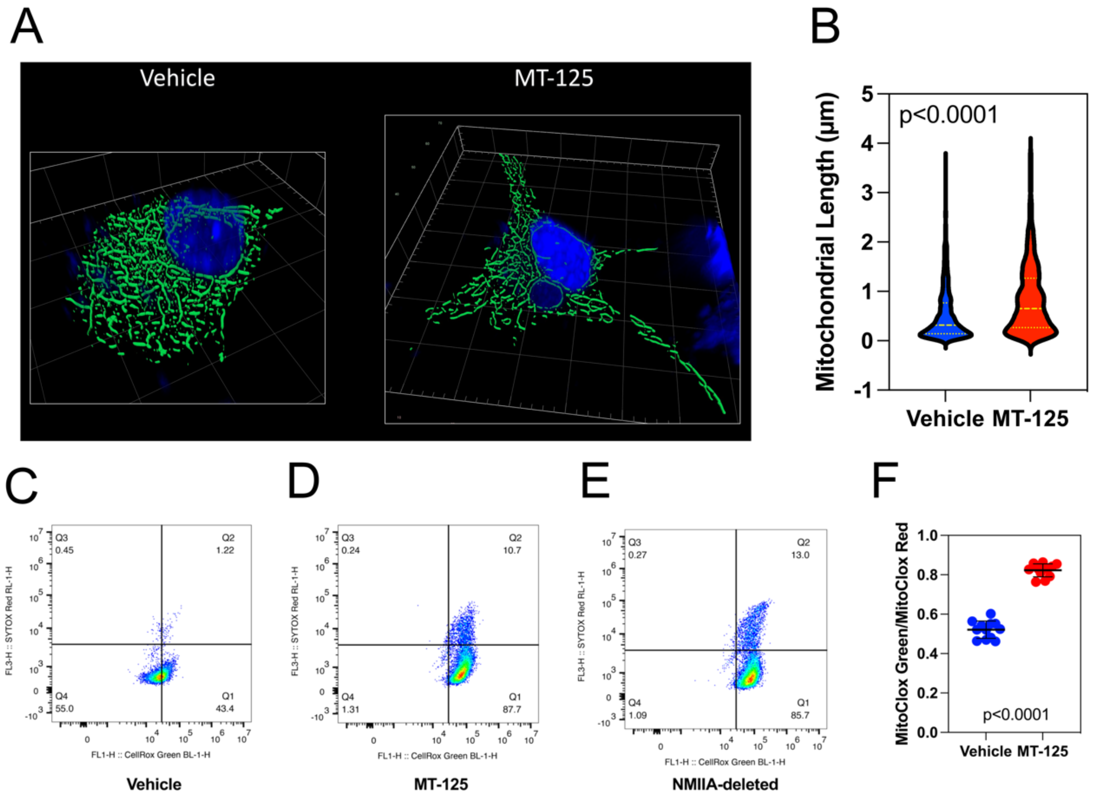

**Figure S4. MT-125 lengthens mitochondria and increases cellular content of reactive oxygen species (ROS).** (**A**). High resolution images of vehicle (*left*) and MT-125 (*right*) treated *Trp53*(-/-) cells stained with DAPI (*blue*) and MitoTracker Green. (**B**). Plot of mitochondrial length for vehicle (*blue*) and MT-125 (*red*) treated cells. Mean length in micrometers ± 1SD are indicated in yellow. (**C-E**): Dual color flow cytometry of *Pten*(-/-) murine GBM cells treated with vehicle (**C**), or MT-125 (**D**) using CellRox Green to measure ROS content and Sytox Red to measure cell viability. Treatment with MT-125 doubles the fraction of cells with detectable ROS staining. (**E**). Similar results are seen in murine GBM cells deleted for *Pten* and NMIIA. (**F**). *Trp53*(-/-) murine GBM cells were treated with vehicle (*blue*) or MT-125 (*red*) and stained with MitoClox—a fluorescent mitochondrial dye with emission shifting from red to green in the presence of mitochondrial ROS. MT-125 increases mitochondrial ROS content by >70%.

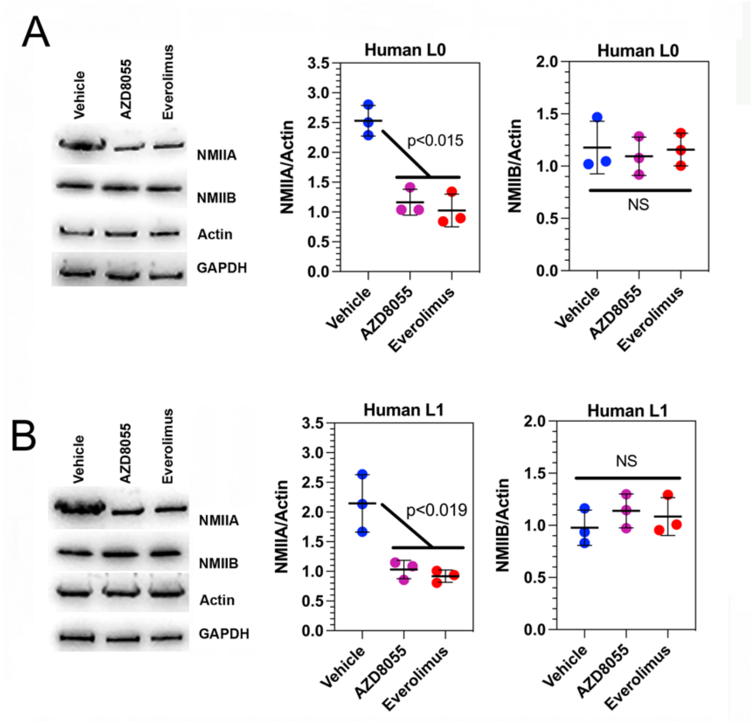

**Figure S5. Inhibition of mTOR decreases expression of NMIIA in two human GBM cell lines**. The human GBM cell lines L0 (**A**) and L1 (**B**) were treated with AZD8055 or everolimus and cell content of NMIIA and IIB were probed on Western blot. Both drugs reduce NMIIA content approximately 2-fold, while having no significant effect on NMIIB content.

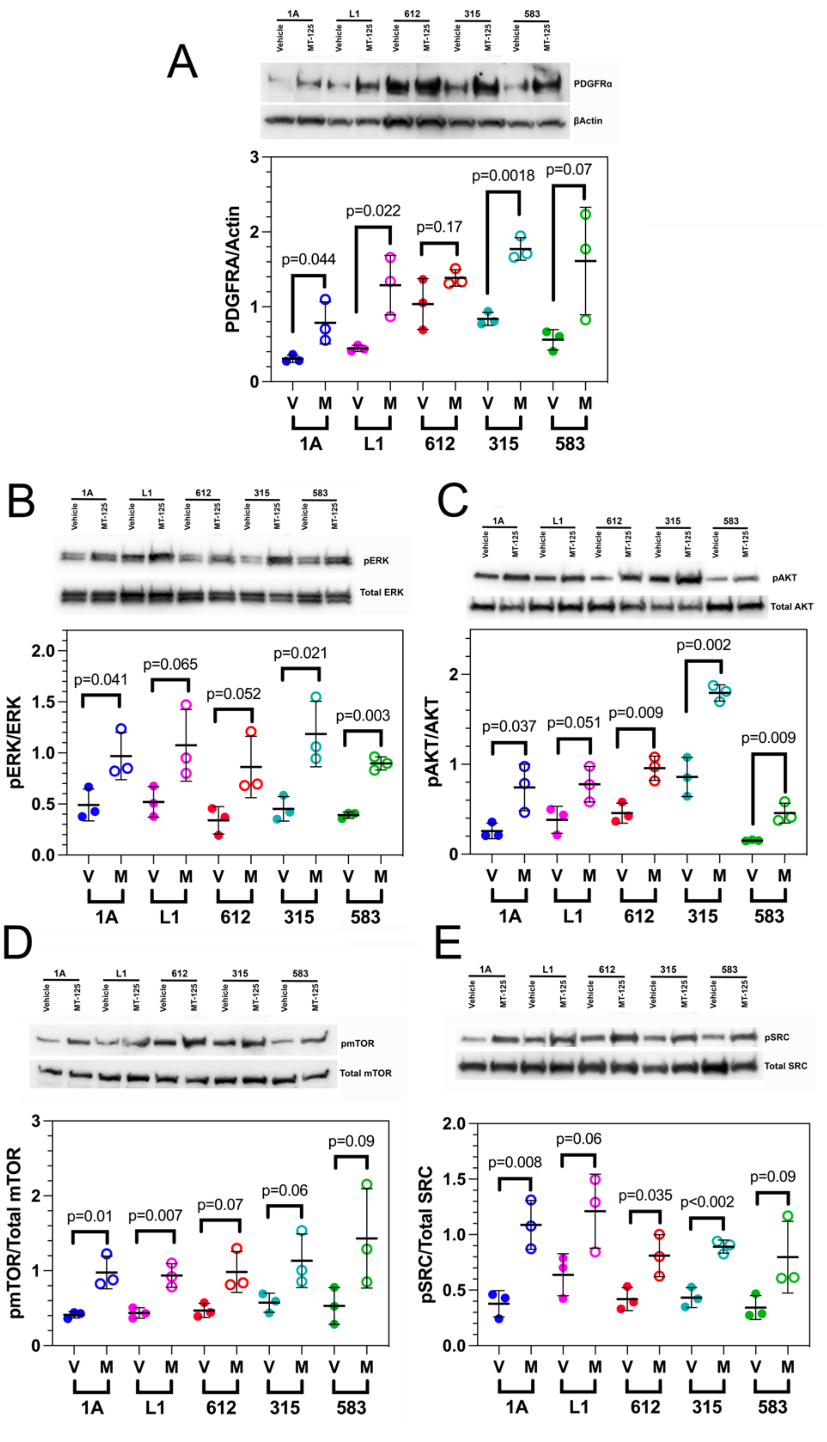

**Figure S6. MT-125 activates PDGFR- and MAPK-dependent signaling in human GBM lines**. Five human GBM cell lines (1A, L1, 612, 315, 583) were treated with MT-125 and cell lysates probed for PDGFRa (**A**), p-ERK1/2 and total ERK1/2 (**B**), p-AKT and total AKT (**C**), p-mTOR and total mTOR (**D**) and p-SRC and total SRC (**E**). In each case, MT-125 significantly increases content of PDGFRa or its downstream activated effectors in a subset of tumor lines.

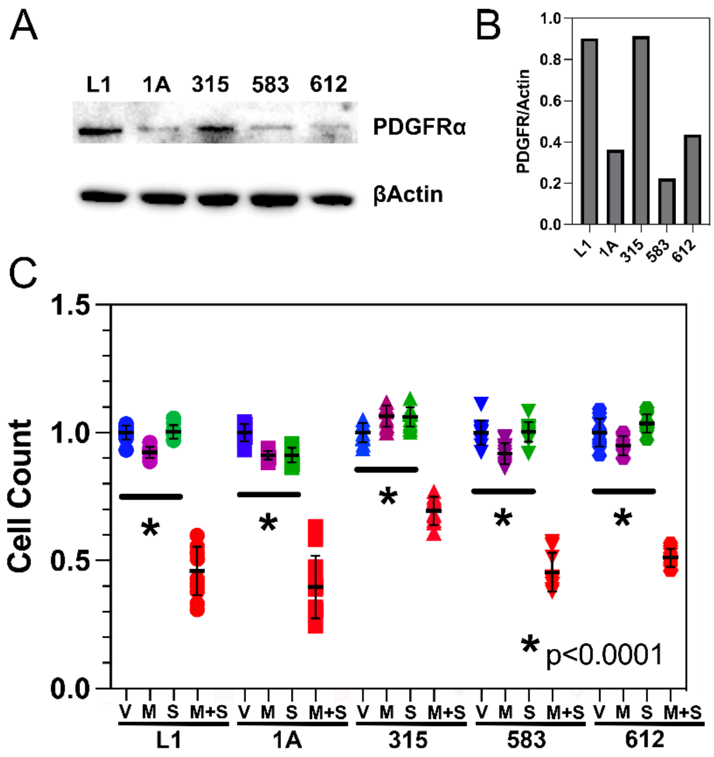

**Figure S7. The effect of combining sunitinib and MT-125 on cell viability in human GBM lines is significantly greater than the effect of either drug alone.** (**A**,**B**). Immunoblots for PDGFRa and Actin (**A**) and plot of normalized PDGFRa immunoblot intensity (**B**) for five primary human GBM lines. Although PDGFRa expression is present in all lines, levels vary across lines. (**C**). These five lines were treated *in vitro* with vehicle (*V, blue*), MT-125 (*M, magenta*), sunitinib (*S, green*), or the combination of MT-125 + sunitinib (*M+S, red*) and cell viability, normalized to vehicle, was measured after 48 hours with CellTiter-Glo. For each line, the combination of MT-125+sunitinib is significantly cytotoxic compared to either drug alone (*p<0.0001, two tailed t test)*.

**KEY RESOURCE TABLE**

| REAGENT or RESOURCE | SOURCE | IDENTIFIER |
|---|---|---|
| Antibodies | | |
| Rabbit polyclonal anti-NMIIA | Biolegend | Cat# 909801; RRID: AB_2565100 |
| Rabbit polyclonal anti-NMIIB | Biolegend | Cat# 909901; RRID: AB_2749903 |
| Mouse monoclonal anti-RAS | Cell Signaling Technology | Cat# 8832; RRID: AB_10978549 |
| Rat monoclonal Anti-HA (clone 3F10) | Sigma-Aldrich | Cat#11867423001 ; RRID: AB_390918 |
| Rabbit anti -PDGFRa | Cell Signaling Technology | Cat# 3164; RRID: AB_2162351 |
| Rabbit monoclonal anti-phospho PDGFRa (pY849) | Cell Signaling Technology | Cat# 3170: RRID: AB_2162348 |
| Rabbit monoclonal anti-AKT | Cell Signaling Technology | Cat# 4691; RRID: AB_915783 |
| Rabbit monoclonal anti-Phospho-AKT (Ser473) | Cell Signaling Technology | Cat# 4060; RRID: AB_2315049 |
| Rabbit anti-ERK | Cell Signaling Technology | Cat# 9102; RRID: AB_330744 |
| Rabbit monoclonal anti-phospho ERK (Thr202/Tyr204) | Cell Signaling Technology | Cat# 4370; RRID: AB_2315112 |
| Rabbit Anti mTOR | Cell Signaling Technology | Cat# 2972; RRID: AB_330978 |
| Rabbit polyclonal anti-phospho-mTOR (Thr2446) | Abcam | Cat#63552; RRID: AB_1140863 |
| Anti IgG Control | Cell Signaling Technology | Cat#3900; RRID: AB_1550038 |
| Anti Integrin β1 | Biolegend | Cat#102211; RRID: AB_492829 |
| Rat monoclonal anti-active integrin β1 | BD Biosciences | Cat#550531; RRID: AB_393729 |
| Rabbit polyclonal anti-Phospho-Src (Tyr418) | Millipore | Cat# 07-909; RRID: AB_568805 |
| Rabbit polyclonal anti-Src | Cell Signaling Technology | Cat# 2108; RRID: AB_331137 |
| Rabbit anti-Phospho-p70 S6 Kinase (Thr389) | Cell Signaling Technology | Cat# 9205; RRID: AB_330944 |
| Rabbit anti-p70 S6 Kinase | Cell Signaling Technology | Cat# 9202; RRID: AB_331676 |
| Mouse monoclonal anti-β-Actin (8H10D10) | Cell Signaling Technology | Cat# 3700; RRID: AB_2242334 |
| Rabbit monoclonal anti-GAPDH | Cell Signaling Technology | Cat# 5174; RRID: AB_10622025 |

| | | |
|---|---|---|
| Goat anti-Rabbit IgG (H+L) Secondary antibody, HRP conjugate | Invitrogen | Cat# 31460 |
| Goat anti-Mouse IgG (H+L) Secondary antibody, HRP conjugate | Invitrogen | Cat# 31430 |
| Bacterial and Virus Strains | | |
| PDGF-IRES-Cre retrovirus | Lei et al., 2011; Kenchappa et al., 2020 | NA |
| Myosin-IIA-GFP plasmid | Addgene | Cat#38297 |
| CMV-GFP-NMHC II-A plasmid | Addgene | Cat#11347 |
| pLV_TurboGFP-Myosin IIA_Mouse-BlastR | This paper | NA |
| pLV_TurboGFP-MYOSIN IIA_Human-BlastR | This paper | NA |
| pLV_TurboGFP-EV | This paper | NA |
| NEB® Stable Competent E. coli | New England BioLabs | Cat#C3040H |
| Biological Samples | | |
| Chemicals, Peptides, and Recombinant Proteins | | |
| PDGF-AA | Peprotech | Cat# 100-13A |
| hFGF (Human Fibroblast Growth factor) | R&D systems | Cat# 233-FB-025 |
| hEGF (Human Epidermal Growth Factor) | Sigma-Aldrich | Cat# E9644 |
| Fibronectin | Millipore-Sigma | Cat# FC010 |
| Heparin | STEMCELL Technologies | Cat# 07980 |
| N2 supplement | Gibco life technologies | Cat# 17502-048 |
| NeuroPlex supplement | Gemini | Cat# 400-161 |
| Rhodamine Phalloidin | Cytoskeleton | Cat# PHDR1 |
| VECTASHIELD with DAPI | Vector Laboratories | Cat# U-1500 |
| Formaldehyde, 10% methanol free | CHEM (VWR) | Cat# 87001-890 |
| SUPER signal West Pico PLUS Chemiluminescent substrate | Thermo Fisher Scientific | Cat# 34580 |
| PBS | Thermo Fisher Scientific | Cat# 21-040 |
| PVDF membrane | Biorad | Cat# 1620174 |
| DMSO | Corning | Cat# 25-950-cqc |
| BSA | Thermo Fisher Scientific | Cat# 23209 |
| Non-fat dry milk (Blotting grade blocker) | Biorad | Cat# 170-6404 |
| Accutase | Sigma-Aldrich | Cat# A6964-500 |
| Ethanol | Thermo Fisher Scientific | Cat# 61500-0020 |
| Tween 20 | Sigma-Aldrich | Cat# P1379 |
| TBS | Thermo Fisher Scientific | Cat# 28358 |
| Protease Inhibitor Cocktail, EDTA-free (100X) | Thermo Fisher Scientific | Cat# 87785 |
| Pierce T-1step transfer buffer | Thermo Fisher Scientific | Cat# 84731 |

| | | |
|---|---|---|
| Western blot striping buffer | Thermo Fisher Scientific | Cat# 46430 |
| 10X Tris/Glycine/SDS buffer | Biorad | Cat# 1610772 |
| Geltrex | Gibco | Cat# A14132-01 |
| Laminin | Thermo Fisher Scientific | Cat# 3400-010-02 |
| RIPA lysis buffer | Thermo Fisher Scientific | Cat# 89900 |
| B27 supplement | Thermo Fisher Scientific | Cat# A3582801 |
| Laemmli SDS Sample Buffer, reducing, 6X | Thermo Fisher Scientific | Cat# AAJ61337AC |
| Antibiotic-Antimycotic (100x) | Gibco | Cat# 15240-062 |
| Blasticidine-S-Hydrochloride | Sigma-Aldrich | Cat# SBR00022 |
| Lenti-X™ Concentrator | Takara | Cat# 631231 |
| Opti-MEM™ I | ThermoFisher | Cat# 31985062 |
| CloneAmp™ HiFi PCR Premix | Takara | Cat# 639298 |
| iScript | BioRad | Cat#1708840 |
| Power SYBR™ Green PCR Master Mix | Thermofisher | Cat#4367659 |
| Lipofectamine™ 3000 | ThermoFisher | Cat# L3000008 |
| High-Capacity Protein A MagBeads | ThermoFisher | Cat#A53035 |
| MitoTracker™ Green | ThermoFisher | Cat#M46750 |
| Hoechst 33342 | ThermoFisher | Cat#62249 |
| Cycloheximide | CellSinaling | Cat#2112S |
| TURBO DNase | ThermoFisher | Cat#AM2238 |
| Superase-IN | ThermoFisher | Cat#AM2694 |
| Anti-HA Magnetic Beads | ThermoFisher | Cat#88836 |
| MitoSOX | ThermoFisher | Cat#M36009 |
| MitoCLox | Lumiprobe | Cat#3549 |
| AZD8055 | Selleck Chemicals | S1555 |
| Everolimus | Selleck Chemicals | S1120 |
| Sunitinib | Selleck Chemicals | S7781 |
| Paxalisib | Selleck Chemicals | S8163 |
| Critical Commercial Assays | | |
| CellTiter-Glo 2.0 assay kit | Promega | Cat# G9242 |
| Pierce™ BCA Protein Assay Kit | Thermo Fisher Scientific | Cat# 23225 |
| RNAqueous™ Total RNA Isolation Kit | Thermo Fisher Scientific | Cat# AM1912 |
| TURBO DNA-free™ Kit | Thermo Fisher Scientific | Cat# AM1907 |
| NEBuilder® HiFi DNA Assembly Master Mix | New England BioLabs | Cat# E2621S |
| PureLink™ DNase Set | ThermoFisher | Cat#12185010 |
| TRIzol™ Plus RNA Purification Kit | ThermoFisher | Cat#A33254 |
| Lenti-X™ p24 Rapid Titer Kit | Takara | Cat#631231 |
| RNeasy Mini kit | Qiagen | Cat# 74104 |
| Active RAS pulldown assay kit | Cell Signaling Technology | Cat#11871/8821 |
| Experimental Models: Cell Lines | | |
| Trp53$^{-/-}$ | This paper | NA |

| | | |
|---|---|---|
| Trp53/Pten$^{-/-}$ | This paper | NA |
| Pten$^{-/-}$ | This paper | NA |
| Pten/NMIIA-/- | This paper | NA |
| Pten/NMIIB-/- | This paper | NA |
| GBM1A | Galli et al., 2004 | NA |
| GBML1 | Deleyrolle et al., 2011 | NA |
| GBML0 | Deleyrolle et al., 2011 | NA |
| GBM612 | Kenchappa et al., 2020 | NA |
| GBM315 | Gift from Dr. Quiñones-Hinojosa | NA |
| GBM509 | Gift from Dr. Quiñones-Hinojosa | NA |
| GBM120 | Gift from Dr. Quiñones-Hinojosa | NA |
| GBM108 | Gift from Dr. Quiñones-Hinojosa | NA |
| GBM593 | Gift from Dr. Quiñones-Hinojosa | NA |
| MES1861 | Reilly et al., 2000; Gursel et al., 2011 | NA |
| Experimental Models: Organisms/Strains | | |
| Trp53$^{fl/fl}$ mice | Jackson Laboratory | Stock# 008462 |
| NOD-SCID mice | Jackson Laboratory | Stock# 005557 |
| Software and Algorithms | | |
| Arivis Vision4D | Zeiss | N/A |
| FIJI | Open source | N/A |
| FlowJo | Becton Dickinson | v10.9 |
| QuantStudio and Analysis | ThermoFisher | v1.42 |

Primers

| Primer | Sequence |
|---|---|
| Myh9 Fw | GGCCCTGCTAGATGAGGAGT |
| Myh9 Rv | CTTGGGCTTCTGGAACTTGG |
| Myh10 Fw | GGAATCCTTTGGAAATGCGAAGA |
| Myh10 Rv | GCCCCAACAATATAGCCAGTTAC |
| Rplp0 Fw | AGATTCGGGATATGCTGTTGGC |
| Rplp0 Rv | TCGGGTCCTAGACCAGTGTTC |
| PDFGRa Fw | GGAGACTCAAGTAACCTTGCAC |
| PDGFRa Rv | TCAGTTCTGACGTTGCTTTCAA |
